# Compartmental Profiling of PDE4B in Systemic Sclerosis

**DOI:** 10.64898/2026.07.27.740454

**Authors:** Astrid Hofman, Pietro Bearzi, Sara Burckhardt, Maryam Asadikorayam, Andrea Laimbacher, Cristian Iperi, Muriel Elhai, Cosimo Bruni, Flurin Struzenegger, Shervin Assassi, Laura Much, Anna-Maria Hoffmann-Vold, Blaž Burja, Helen C. Jarnigan, Michael L. Whitfield, Mike O. Becker, Lumeng Li, Philip Stauffer, Shihan Xu, Sophie Wagner, Yasmine Illi, Zhiyun Gong, Elena Pachera, Oliver Distler

## Abstract

**Objectives:** The preferential phosphodiesterase 4B (PDE4B) inhibitor nerandomilast was recently approved for treatment of idiopathic pulmonary fibrosis (IPF) and progressive pulmonary fibrosis. Its proposed immunomodulatory, anti-fibrotic, and endothelial-stabilising actions target all three cardinal features of SSc, yet PDE4B expression has not been systematically characterised in SSc tissue. We aimed to define PDE4B expression across fibrotic organs and cellular compartments in SSc.

**Methods:** PDE4B expression was profiled in SSc lung, peripheral blood mononuclear cells (PBMCs) and skin on the transcript level using single-cell RNA sequencing data and on the protein level using immunohistochemistry, immunofluorescence and multiplexed immunofluorescent stainings.

**Results:** PDE4B was consistently dysregulated in immune cells across SSc tissue and PBMCs, with compartment-specific direction and distribution. In SSc-ILD lung, expression was increased in CD8⁺ and CD4⁺ memory T-cells. In PBMCs, expression was increased in B cells, monocytes, and CD8⁺ T-cells, and stratified patients into three endotypes (PDE4B^ˡᵒ^/^ᵐᵉᵈ^/^hi^) not distinguishable by clinical variables. In skin, bulk RNA-seq showed a significant global increase, which localized to myeloid cells in scRNA-seq data. Approximately 90% of FAP⁺ activated fibroblasts co-expressed PDE4B at the protein level in SSc skin, identifying the activated fibroblast compartment as a candidate target for PDE4B inhibition. No PDE4B dysregulation was detected in vascular cell types.

**Conclusions:** This first cell-type-resolved characterisation of PDE4B in SSc demonstrates consistent immune-cell dysregulation across tissues and protein-level enrichment in activated fibroblasts. This provides a human-tissue rationale for the immunomodulatory and anti-fibrotic effects of PDE4B inhibition and supporting PDE4B as a disease-relevant therapeutic target in SSc.

**Key messages:** *What is already known on this topic:*

- Nerandomilast (BI 1015550), a PDE4B-preferential inhibitor, was approved for idiopathic pulmonary fibrosis and progressive pulmonary fibrosis.
- Pre-clinical studies indicate that PDE4B inhibition may act on all cardinal features of SSc.

*What this study adds:*

- First cell-type-resolved characterization of PDE4B expression across SSc-affected lung, PBMCs, and skin.
- PBMC PDE4B expression is heterogeneous, stratifying patients into PDE4Bˡᵒ/ᵐᵉᵈ/ʰⁱ endotypes independent of standard clinical variables.
- scRNA-seq shows increased myeloid PDE4B expression in SSc skin, while ∼90% of FAP⁺ activated fibroblasts in SSc skin express PDE4B protein.

*How this study might affect research, practice or policy:*

- The study strengthens the human-level evidence underpinning the target rationale for PDE4B inhibition in SSc.

## Introduction

Systemic sclerosis (SSc) is a complex autoimmune rheumatic disease characterized by a triad of manifestations: microvasculopathy, autoimmunity/inflammation, and fibrosis.^1,2^ The variability in severity and type of organ involvement result in a heterogeneous prognosis, which makes clinical management and trial design challenging.^3^ As highlighted in the 2023 updated EULAR treatment recommendations and the ERS/EULAR guidelines for the management of connective tissue disease-associated interstitial lung diseases (CTD-ILD), several pharmacological treatments are available for SSc.^3,4^ Established treatment strategies for SSc include immunosuppressive treatments such as cyclophosphamide, mycophenolate mofetil (MMF), and methotrexate, depending on the underlying disease manifestations.^5,6^ Tocilizumab, an anti-interleukin (IL)-6 receptor antibody, has shown efficacy for SSc-ILD in patients with early, inflammatory diffuse cutaneous (dc)SSc.^3,7^ Rituximab, an anti-CD20 antibody, approved in Japan, has also been included in the recommendations for both SSc skin fibrosis and SSc-ILD.^3^ Nintedanib has additionally been shown to be effective in treating SSc-ILD.^8,9^ While these therapies may slow the progression of fibrosis, they fail to reverse or stop it, and are not disease modifying.^10^ Thus, there remains an unmet need for therapies capable of simultaneously addressing the vascular, inflammatory, and fibrotic components of SSc.

Recently, Nerandomilast (BI 1015550), an orally administered selective phosphodiesterase (PDE)4B inhibitor was approved by the Food and Drug Administration (FDA) for the treatment of idiopathic pulmonary fibrosis (IPF) and progressive pulmonary fibrosis (PPF) of different origins after meeting the primary endpoint in two Phase III clinical trials.^11^ In addition to slowing forced vital capacity (FVC) decline, it shows a morality benefit, meeting a major unmet need. This implies an effect on overall disease progression, consistent in the subgroup of patients with autoimmune ILDs, including SSc-ILD.^12^ PDE4B-targeting with Nerandomilast is currently being tested in several randomized controlled trials (RCTs), including a placebo-controlled, multicenter, international RCT aiming to test disease modification in SSc (NCT07497087).

PDEs are enzymes encoded by 21 genes and divided into 11 families based on their structural similarity.^13^ PDEs mediate hydrolysis of cyclic adenosine monophosphate (cAMP) and cyclic guanosine monophosphate (cGMP) to 5’-AMP and 5’-GMP, respectively.^10,13^ These are second messengers that activate protein kinases, which in turn regulate numerous downstream targets, such as receptors, ion channels, cytoskeletal proteins and transcription factors.^14^ The PDE4 family comprises four subtypes: PDE4A, B, C, and D, all selectively hydrolyzing cAMP.^10,14^ Of these, PDE4B is the most widely expressed, specifically in immune cells and the lungs.^14^ Inhibition of PDE4B prevents hydrolysis of cAMP, thereby increasing intracellular cAMP levels which decreases the inflammatory response.^15^

PDE4B inhibition has been proposed to act on each of the three characterizing features of SSc. Regarding vasculopathy, PDE4B inhibition was found to strengthen endothelial junctions via inhibition of adhesion proteins in human lung microvascular endothelial cells.^16^ With respect to inflammation, nerandomilast was found to suppress lipopolysaccharide (LPS)-induced tumour necrosis factor (TNF)-α and Interleukin (IL)-2 synthesis in human peripheral blood mononuclear cells (PBMCs).^17^ In a bleomycin-induced SSc-ILD model, nerandomilast reduced the inflammatory cell influx in bronchoalveolar lavage (BAL) fluid, along with suppressing multiple pro-inflammatory cytokines. Furthermore, the number of infiltrating CD3^+^ T-cells and F4/80^+^ macrophages in fibrotic lung tissue was reduced in the same study.^15^ Concerning fibrosis, cAMP accumulation inhibited myofibroblast transformation and attenuated TGF-β-mediated contractility.^16^ Nerandomilast was additionally found to alleviate skin fibrosis in a bleomycin mouse model, accompanied by a decrease in fibrosis-related genes and proteins.^15^

Taken together, the proposed immunomodulatory, direct anti-fibrotic, and endothelial-stabilizing properties of PDE4B align with the key pathogenic features of SSc, suggesting a mechanistic rationale that may extend beyond existing therapies. However, PDE4B expression has not been systematically characterized in human SSc tissues. Since increased expression of PDE4B in the affected tissue is central to the therapeutic rationale, this represents a critical knowledge gap, in addition to helping to better understand heterogeneity that may underline (non-)response. Therefore, this study aims to investigate whether PDE4B expression is consistently dysregulated across fibrotic organs and cellular compartments in SSc, supporting its potential as a disease-relevant target for pharmacological intervention.

## Results

### PDE4B expression is upregulated in CD8^+^ T-cells and CD4^+^ memory T-cells in SSc-ILD

We started by analyzing lung tissue, where RCTs have established clinical efficacy of PDE4B inhibition in fibrotic lung diseases. Expression of PDE4B in SSc-ILD and IPF lungs therefore provides a validated benchmark for a viable clinical target. All lung samples were obtained from end-stage patients (both SSc-ILD and IPF) undergoing lung transplantation surgery, whereas healthy lung tissue was obtained following rejection as candidate donors for transplant (**Table 1**). Histopathological information can be found in the respective studies. Two comparisons were performed for the scRNA-seq analysis for the lung, PBMCs and skin. This included a “single-cell” analysis, in which each cell was treated as an independent observation, and a “(patient) pseudobulk” analysis, in which PDE4B expression was averaged across all cells per donor before group comparison.

**Table 1.** Overview of Datasets used.

| **Tissue** | **Number of Patients** | **Dataset Type** | **Dataset Source** | **Publication** |
| --- | --- | --- | --- | --- |
| **Lung** | IPF lung: n = 5<br>Healthy lung: n = 4 | scRNA-seq | GSE128033<sup>32</sup> | Morse et al. (2019); doi: 10.1183/13993003.02441-2018. |
| **Lung** | SSc-ILD: n = 4 | scRNA-seq | GSE128169<sup>33</sup> | Valenzi et al (2019); doi: 10.1136/annrheumdis-2018-214865 |
| **Peripheral Blood Mononuclear Cells (PBMCs)** | SSc: n = 25<br>Healthy: n = 9 | scRNA-seq | In-house | Present study |
| **Skin** | SSc: n = 48<br>Healthy: n = 33 | Bulk RNA-seq | PRESS cohort<sup>18</sup> | Skaug et al. (2020); doi: 10.1136/annrheumdis-2019-215894. |
| **Skin** | SSc: n = 30 | scRNA-seq | GSE209635<sup>20</sup> | Khanna et al (2022); doi: 10.1172/jci.insight.159566 |
| **Skin** | SSc: n = 12<br>Healthy: n = 10 | scRNA-seq | GSE138669<sup>19</sup> | Tabib et al (2021); doi: 10.1038/s41467-021-24607-6 |
| **Skin** | SSc: n = 13<br>Healthy: n = 10 | Immunohisto-chemical stainings | In-house | Present study |
| **Skin** | SSc: n = 5<br>Healthy: n = 3 | Immunofluorescent stainings | In-house | Present study |

The scRNA-seq of lung biopsies from patients with SSc-ILD, IPF and HCs revealed 17 distinct cell populations (53,471 cells) annotated with top markers that recapitulate the known cell types in the lung (**Figure 1A, Figure S1A**). PDE4B expression was highest in the immune cell populations, namely both CD8^+^ and CD4^+^ T-cells, macrophages, B-cells, plasma cells, dendritic cells (DCs), NK cells, and endothelial cells (**Figure 1B**). When comparing the average PDE4B expression across all cell types, both CD8^+^ and CD4^+^ T-cells had the strongest increase in expression in SSc-ILD and IPF relative to HCs. Conversely, macrophages and DCs showed a decreased average expression of PDE4B in SSc-ILD and IPF compared to HCs (**Figure 1A C**). Macrophages and DCs showed a significant decrease in PDE4B expression in IPF relative to HCs not observed in SSc-ILD versus HCs **(Figure S1B-C)**. In contrast, CD8^+^ T-cells showed an increased expression in SSc-ILD as compared to HCs not detected in IPF relative to HCs **(Figure S1B-C).** CD4^+^ T-cells showed no significant change in PDE4B expression in patient-level differences in the disease groups compared to HCs **(Figure S1B-C).** This indicates that alterations of PDE4B expression in the fibrotic lung are primarily attributed to immune cells, with a low average transcript detection in fibroblasts and endothelial cells.

**Figure 1.**
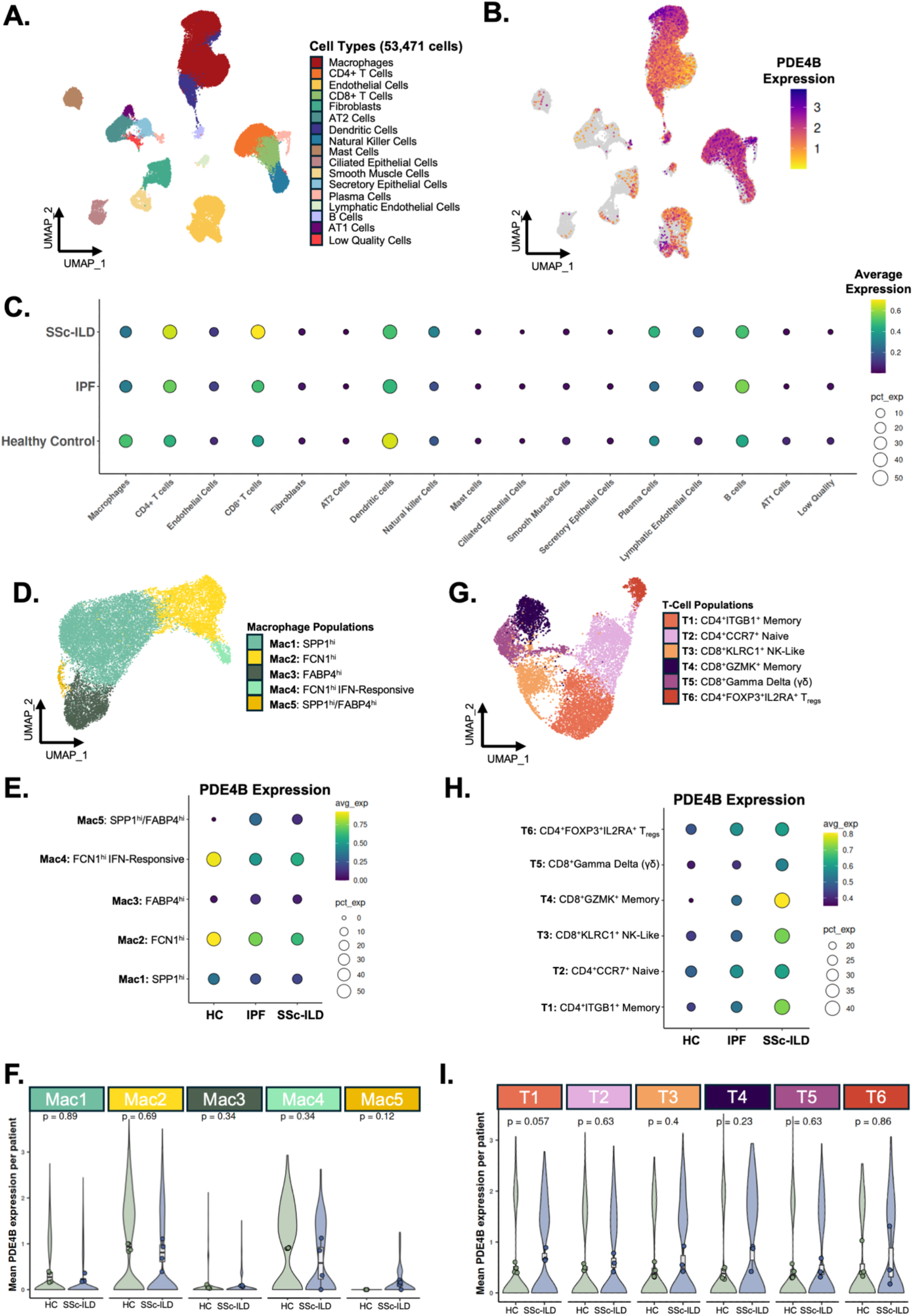
scRNA-seq of lung biopsies in SSc-ILD and IPF reveals an upregulation of PDE4B expression in CD8+ T-cells and a decreased expression in macrophages. (A) UMAP showing integrated scRNA-seq data (53,471 cells) from GSE128169 and GSE128033, from SSc-ILD (n=4), IPF (n=5) and HCs (n=4). Annotated cell types include macrophages, CD4^+^ T-cells, CD8^+^ T-cells, dendritic cells, natural killer (NK) cells, mast cells, B-cells, plasma cells, endothelial cells (ECs), lymphatic ECs, smooth muscle cells, fibroblasts, alveolar type (AT)1 and AT2 cells, ciliated epithelial cells, and a cluster of low-quality cells. (B) UMAP showing cell-type expression of PDE4B, which localizes to macrophages, B-cells, NK cells, CD8^+^ and CD4^+^ T-cells and dendritic cells. (C) Dot plot of PDE4B expression across cell populations in the lung for HCs, IPF and SSc-ILD, with dot size representing the percentage of expressing cells and colour indicating average expression level. (D) UMAP of macrophage subpopulations (25,815 cells) coloured by population. (E) Dot plot of PDE4B expression across macrophage subpopulations in HC, IPF and SSc-ILD, with dot size representing the percentage of expressing cells and colour indicating average expression level. (F) Mean PDE4B expression per patient across macrophage subpopulations in HCs and SSc-ILD. Statistical comparisons were performed using the Wilcoxon rank-sum test. (G) UMAP of the T-cell populations coloured by population. (H) Dot plot of PDE4B expression across T-cell subpopulations in HC, IPF and SSc-ILD, with dot size representing the percentage of expressing cells and colour indicating average expression level. (I) Mean PDE4B expression per patient across T-cell subpopulations in HC and SSc-ILD. Statistical comparisons were performed using the Wilcoxon rank-sum test. Abbreviations: scRNA-seq: single cell RNA sequencing, HCs: healthy controls, IPF: diopathic pulmonary fibrosis, SSc-ILD: systemic sclerosis-associated interstitial lung disease

To further characterize the PDE4B expression patterns within immune compartments, we examined their subpopulations. To this end, the macrophages were subclustered into five distinct subsets (**Figure 1D, Figure S2A**). Among these, the FCN1^hi^ monocyte-derived subsets (Mac2 and Mac4) showed the highest PDE4B expression in HCs, with low expression across other subpopulations (**Figure 1E**). However, when comparing mean PDE4B expression across patients, no macrophage subpopulation shows a significant difference in SSc-ILD versus HCs (**Figure 1F**). Conversely, in IPF, the Mac4 population exhibited a significant decrease in PDE4B expression, while the Mac3 and Mac5 populations show a significant increase compared to HCs, despite a low overall expression level **(Figure S2B).**

To understand how T-cells were generally altered in their PDE4B expression profiles, both CD8^+^ T-cells and CD4^+^ T-cells were subclustered together and divided into six populations based on their top markers (**Figure 1G, Figure S2C**). Across the T-cell subpopulations, PDE4B expression followed a disease-associated gradient: highest in SSc-ILD, intermediate in IPF, and lowest in HCs. More specifically, the T1, T3, and T4 populations had the highest average expression in SSc-ILD **(Figure 1H)**. Furthermore, the T1 (CD4^+^ITGB1^+^ Memory) population demonstrated the largest increase in PDE4B expression when comparing SSc-ILD to HCs **(Figure 1I)**. Interestingly, no T-cell populations significantly differed in their PDE4B expression between IPF and healthy controls **(Figure S2D)**. The elevated PDE4B expression in SSc-ILD CD8^+^ T-cells was not attributable to any single subpopulation, suggesting a pan-CD8^+^ shift rather than subset-specific upregulation. In contrast, the CD4^+^ compartment showed no global difference, with the increase restricted to the memory subpopulation in SSc-ILD. Altogether, SSc-ILD shows a distinct PDE4B signature suggesting disease-specific immune mechanisms.

### Heterogeneous PDE4B expression patterns across SSc PBMC populations identify a high-expressing patient endotype

Next, we examined PBMCs, an accessible biomaterial well suited to precision medicine applications. ScRNA-seq was performed on PBMCs from SSc patients and HCs (**Supplemental File 1, Table 2)**. In short, SSc patients all met the 2013 ACR/EULAR criteria for SSc diagnosis, with healthy controls being age-and sex-matched. SSc patients were early, active patients that were either non-responsive to the first line of therapy or treatment naïve but requiring the start of immunosuppressive treatment.

**Table 2.** Patient characteristics for the PBMC scRNA-seq cohort.

| **<u>Characteristic</u>** | **<u>SSc (n = 25)</u>** | **<u>Healthy Controls (n = 9)</u>** |
| --- | --- | --- |
| **Age, years, median [Q1-Q3]** | 52 [49-62] | 56 [48-59] |
| **Sex - Female n (%)** | 14 (56%) | 8 (89%) |
| **<u>Disease Duration</u>** |  |  |
| **Time from first non-Raynaud's manifestation, months, median [Q1-Q3]** | 12 [18 - 54] |  |
| **<u>Autoantibodies</u>** |  |  |
| **Anti-Scl70, n (%)** | 17 (68%) |  |
| **ACA, n (%)** | 3 (12%) |  |
| **RNPIII, n (%)** | 3 (12%) |  |
| **<u>Disease Characteristics</u>** |  |  |
| **dcSSc, n (%)** | 9 (36%) |  |
| **mRSS, median [Q1-Q3]** | 8 [0 - 17] |  |
| **ILD, n (%)** | 22 (88%) |  |
| **FVC, %, median [Q1-Q3]** | 76% [71-92] |  |
| **DLCO %, median [Q1-Q3]** | 62% [47-76] |  |
| **KCO (DLCO/VA) %, median [Q1-Q3]** | 80% [63-85] |  |
| **CRP, median [Q1-Q3]** | 0.57 mg/dL [0.15-1.1] |  |
| **ESR, median [Q1-Q3]** | 23.5 mm/h [9.5-32.25] |  |
| **Disease Activity: rEUSTAR-AI, median [Q1-Q3]** | 3.25 [1.67 - 5.00] |  |
| **Naive to immunosuppressive therapy, n (%)** | 20 (80%) |  |

Overall, 108,138 cells were automatically annotated to the major cell populations in PBMCs, where PDE4B was found to be broadly expressed across cell types (**Figure 2A-B**). On a cellular level, PDE4B expression was significantly increased in B-cells, monocytes and CD8^+^ T-cells in SSc patients compared to HC, identifying these compartments as priorities for higher-resolution analysis (**Figure 2C**). When comparing mean patient-level PDE4B expression across SSc patients and HCs, no cell population demonstrated a significant increase in expression. However, B-cells had the largest increase across all cell types **(Figure S3A).**

**Figure 2.**
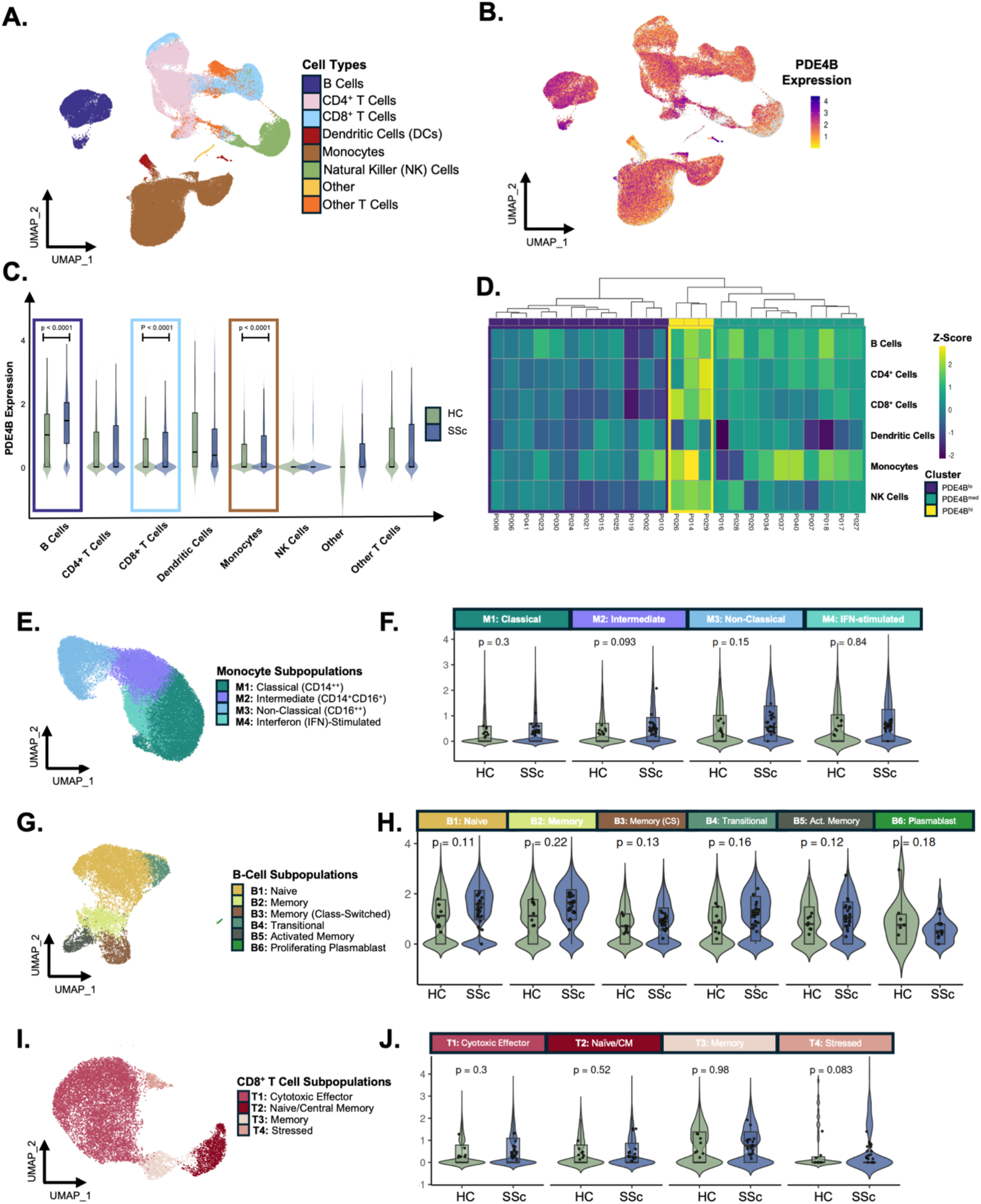
PDE4B expression is upregulated in non-classical monocytes, activated memory B-cells and CD8^+^ T-cells in PBMCs from SSc patients. **(A)** UMAP showing integrated data of 108,138 PBMCs from SSc patients (n = 25) and HCs (n = 9), coloured by annotated cell type. **(B)** UMAP showing the PDE4B expression levels across cell types as defined in (A). **(C)** Violin plots showing PDE4B expression across major PBMC cell types in healthy controls and SSc patients. Statistics performed with the *MAST* framework, adjusted p-value labelled in figure. **(D)** Heatmap displaying hierarchical clustering of PDE4B Z-scores across major PBMC cell types per patient, normalised to the mean expression of healthy controls. Three patient clusters with distinct PDE4B expression levels are identified: PDE4B^lo^, PDE4B^med^, and PDE4B^hi^. **(E)** UMAP of the monocyte subpopulations coloured by subpopulation. **(F)** Violin plots showing mean PDE4B expression across monocyte subpopulations in healthy controls and SSc patients, where each dot is an individual sample. Statistical comparisons were performed using the Wilcoxon rank-sum test. **(G)** UMAP of the B-cell subpopulations coloured by subpopulation. **(H)** Violin plots showing mean PDE4B expression across B-cell subpopulations in healthy controls and SSc patients, where each dot is an individual sample. Statistical comparisons were performed using the Wilcoxon rank-sum test. **(I)** UMAP of the CD8^+^ T-cell subcluster coloured by subpopulation. **(J)** Violin plots showing mean PDE4B expression across CD8^+^ T-cell subpopulations in healthy controls and SSc patients, where each dot is an individual sample. Statistical comparisons were performed using the Wilcoxon rank-sum test. Abbreviations: SSc: Systemic sclerosis; HC: Healthy controls.

**Table 3.**
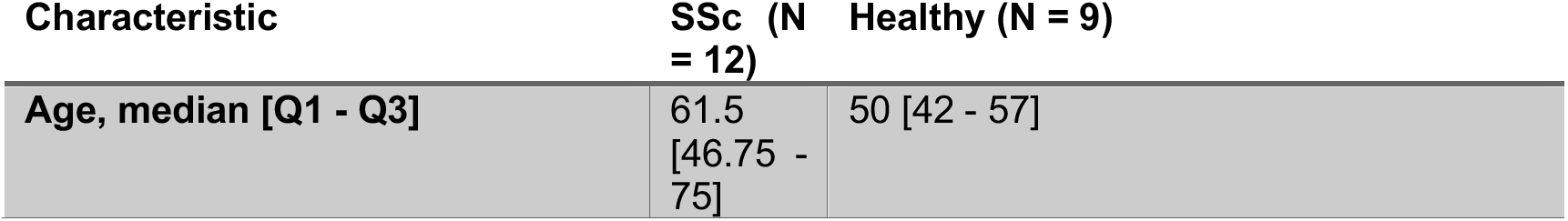
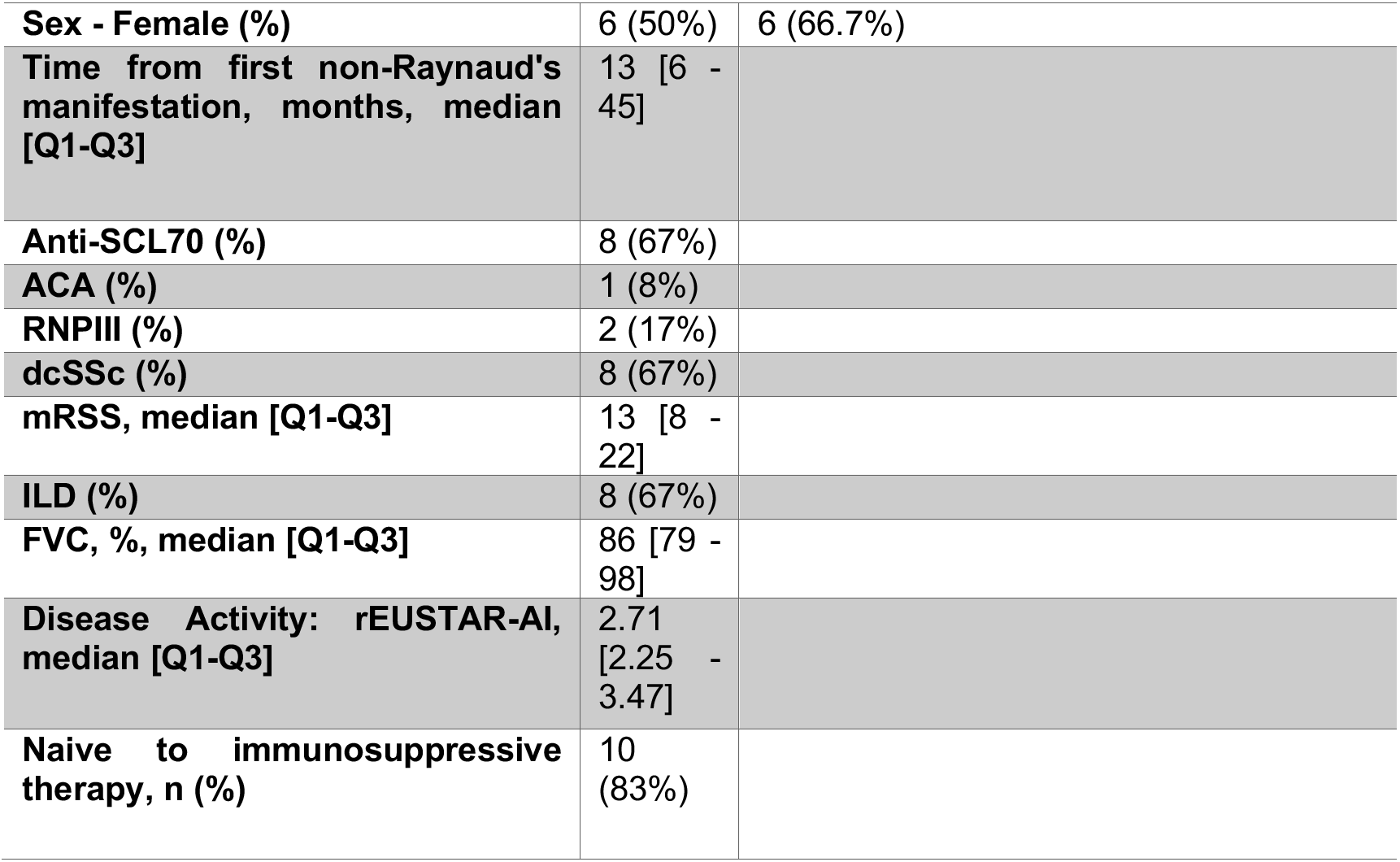
Patient demographics for the IHC cohort.

| Characteristic | SSc (N = 12) | Healthy (N = 9) |
| --- | --- | --- |
| Age, median [Q1 - Q3] | 61.5 [46.75 - 75] | 50 [42 - 57] |
| **Sex - Female (%)** | 6 (50%) | 6 (66.7%) |
| **Time from first non-Raynaud's manifestation, months, median [Q1-Q3]** | 13 [6 - 45] |  |
| **Anti-SCL70 (%)** | 8 (67%) |  |
| **ACA (%)** | 1 (8%) |  |
| **RNPIII (%)** | 2 (17%) |  |
| **dcSSc (%)** | 8 (67%) |  |
| **mRSS, median [Q1-Q3]** | 13 [8 - 22] |  |
| **ILD (%)** | 8 (67%) |  |
| **FVC, %, median [Q1-Q3]** | 86 [79 - 98] |  |
| **Disease Activity: rEUSTAR-AI, median [Q1-Q3]** | 2.71 [2.25 - 3.47] |  |
| **Naive to immunosuppressive therapy, n (%)** | 10 (83%) |  |

**Table 4.**
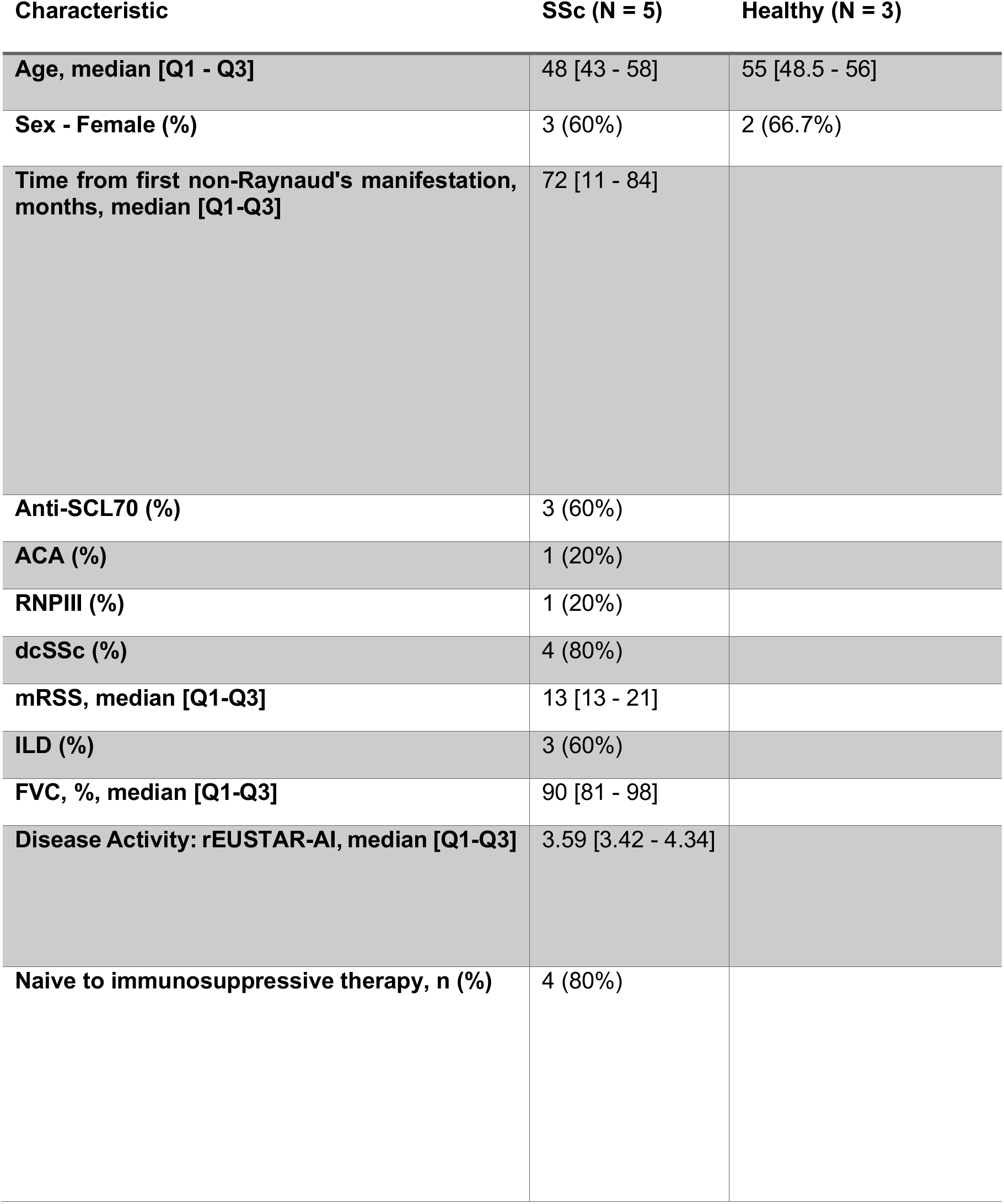
Patient demographics for the IF cohort.

| **Characteristic** | **SSc (N = 5)** | **Healthy (N = 3)** |
| --- | --- | --- |
| **Age, median [Q1 - Q3]** | 48 [43 - 58] | 55 [48.5 - 56] |
| **Sex - Female (%)** | 3 (60%) | 2 (66.7%) |
| **Time from first non-Raynaud's manifestation, months, median [Q1-Q3]** | 72 [11 - 84] |  |
| **Anti-SCL70 (%)** | 3 (60%) |  |
| **ACA (%)** | 1 (20%) |  |
| **RNPIII (%)** | 1 (20%) |  |
| **dcSSc (%)** | 4 (80%) |  |
| **mRSS, median [Q1-Q3]** | 13 [13 - 21] |  |
| **ILD (%)** | 3 (60%) |  |
| **FVC, %, median [Q1-Q3]** | 90 [81 - 98] |  |
| **Disease Activity: rEUSTAR-AI, median [Q1-Q3]** | 3.59 [3.42 - 4.34] |  |
| **Naive to immunosuppressive therapy, n (%)** | 4 (80%) |  |

**Table 5.** Patient demographics for the Multiplex-IF cohort.

| **Characteristic** | **SSc (n = 3)** | **HC (n = 1)** |
| --- | --- | --- |
| **Age, median [Q1 - Q3]** | 65 [60.5 - 67] | 40 |
| **Sex - Female (%)** | 1 (33%) | 1 (100%) |
| **Time from first non-Raynaud's manifestation, months, median [Q1-Q3]** | 12 [10 -36] |  |
| **Anti-SCL70 (%)** | 1 (33%) |  |
| **ACA (%)** | 1 (33%) |  |
| **RNPIII (%)** | 0 (0%) |  |
| **dcSSc (%)** | 3 (100%) |  |
| **mRSS, median [Q1-Q3]** | 28 [19 -32] |  |
| **ILD (%)** | 2 (67%) |  |
| **FVC, %, median [Q1-Q3]** | 81 [75 - 81] |  |
| **Disease Activity: rEUSTAR-AI, median [Q1-Q3]** | 4.75 [4.42 - 4.75] |  |
| **Naive to immunosuppressive therapy, n (%)** | 3 (100%) |  |

Although differences were apparent at the cellular level, they did not translate to patient-level differences, prompting us to examine the heterogeneity of PDE4B expression across patients and HCs. Unsupervised clustering of PDE4B Z-scores did not separate patients from HCs, indicating that PDE4B expression alone does not distinguish SSc from HCs at the population level **(Figure S3B)**. This observation could be due to heterogeneity in the SSc group, with only a proportion of patients having increased PDE4B levels. We therefore computed PDE4B Z-scores for patients relative to the mean HC expression, which stratified them into three distinct populations: PDE4B^lo^, PDE4B^med^, and PDE4B^hi^ (**Figure 2D**). Notably, the PDE4B^hi^ group showed elevated expression across multiple cell types, suggesting a PDE4B^hi^ endotype that may be particularly amenable to therapeutic PDE4B inhibition.

Given the identification of a PDE4B^hi^ subgroup, we investigated whether the level of PDE4B expression in PBMCs associates with specific clinical characteristics, which could inform patient stratification in future clinical trials. However, none of the clinical parameters tested were significantly associated to PDE4B expression levels **(Figure S3C)**. This lack of correlation suggests that PDE4B expression in PBMCs may be regulated independently of the tested clinical variables and that direct biomarker measurements instead of clinical parameters are needed to enrich patients with PDE4B^hi^ expression for clinical trials. Notably, it was found that immunosuppression did not influence PDE4B expression patterns across the endotypes (p = 0.82) indicating that PDE4B might not be targeted by standard background treatments.

Next, we asked whether PDE4B expression is altered in specific cell subsets within the monocyte, B-cell and CD8^+^ T-cell compartments. Monocytes resolved into four distinct subpopulations based on established transcriptional and cell-surface markers (**Figure 2E, Figure S4A-C)**. Non-classical monocytes showed the highest average PDE4B expression in SSc patients on the cell-level, whereas comparing mean expression across patients showed a disease-associated increase in intermediate monocytes (**Figure 2E-F, Figure S4D)**. B-cells (12,645 cells) partitioned into six subsets, where naïve, memory, and transitional B-cells showed an increased average PDE4B expression in SSc that did not translate to significant changes when comparing patient mean expression (**Figure 2G-H**, **S4E-H**). Lastly, CD8⁺ T-cells comprised four distinct subsets, where increased PDE4B average expression was primarily observed in the CD8^+^ memory subset (**Figure 2I**, **Figures S4I-L**). On the patient-level, PDE4B expression was primarily increased in the stressed CD8^+^ T-cells of SSc patients compared to HCs (**Figure 2J**). Overall, SSc-associated patterns of PDE4B expression were observed across monocyte, B-cell, and CD8^+^ T-cell subsets. These changes were not uniformly reflected at the patient level, consistent with the PDE4B expression endotype stratification.

### PDE4B expression in SSc skin is primarily observed in myeloid cell populations and FAP^+^ cells

Finally, we aimed to identify whether PDE4B was dysregulated in SSc skin. First, we utilized bulk RNA-seq from the Prospective Registry of Early Systemic Sclerosis (PRESS) to assess global expression of PDE4B in a large cohort of skin biopsies from early dcSSc patients (n=48) and HCs (n=33). Patients had a disease duration < 3 years with 60.5% having immunosuppressive treatment. HCs were age-, sex-and ethnicity-matched.^18^ Normalized expression of RNA-seq data demonstrated a significant global increase of PDE4B expression in SSc patients compared HCs (Log2FC = 1.35, p < 0.00001, **Figure 3A**).

**Figure 3.**
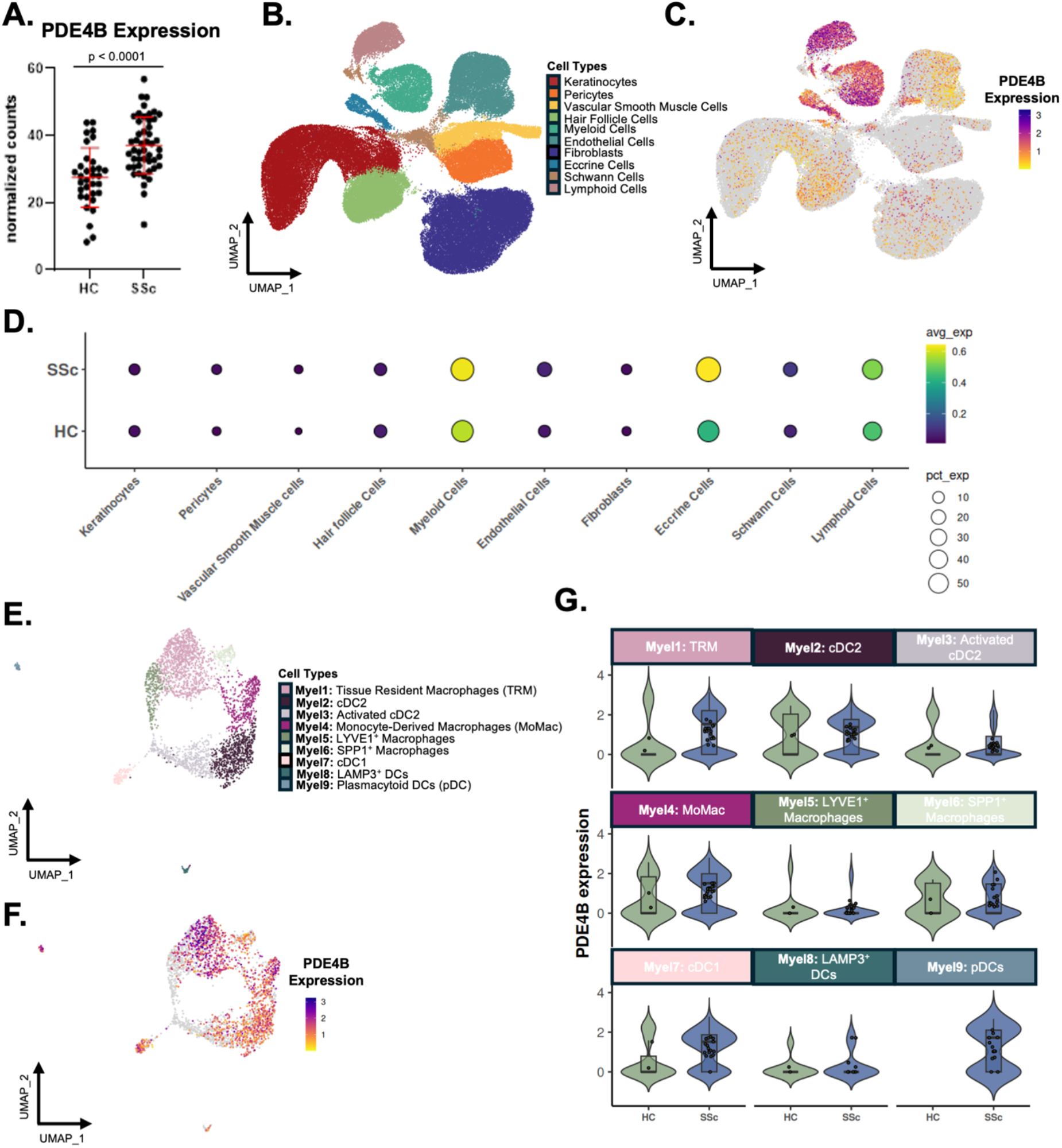
PDE4B expression in SSc skin is primarily observed in myeloid cells and perivascular regions. **(A)** Normalised PDE4B transcript counts from bulk RNA-seq of the PRESS cohort, comparing SSc patients (n = 48) and age-and sex-matched healthy controls (HC, n = 33). PDE4B expression was significantly increased in SSc compared to HC (log2FC = 1.35, p < 0.0001, t-test). **(B)** UMAP of 91,196 cells of SSc patient and healthy control skin, coloured by annotated cell type. This revealed ten populations: keratinocytes, pericytes, vascular smooth muscle cells, hair follicle cells, myeloid cells, endothelial cells, fibroblasts, eccrine cells, Schwann cells, and lymphoid cells. **(C)** Feature plot showing PDE4B expression across cell types in both healthy and SSc skin. **(D)** Dot plot showing the raw PDE4B expression across the different cell types as annotated in (B). Dot colouring indicates the average expression of PDE4B, whereas the dot size indicates the percentage of cells expressing PDE4B, split by SSc skin and HCs. **(E)** UMAP of integrated data showing the nine identified myeloid cell populations: tissue-resident macrophages, cDC2, activated cDC2, monocyte-derived macrophages, LYVE1^+^ macrophages, SPP1^+^ macrophages, cDC1, LAMP3^+^ DCs, plasmacytoid DCs. **(F)** Feature plot showing the expression of PDE4B across myeloid cell populations. **(G)** Patient mean expression of PDE4B plotted for each myeloid cell subpopulation, where each dot represents one patient (n = 2-19). Abbreviations: SSc: Systemic sclerosis; HC: Healthy controls; DC: dendritic cells.

To identify cell type-specific changes in expression, scRNA-sequencing data from SSc and HC skin were used from two publicly available datasets.^19,20^ All patients had dcSSc, with a short disease duration (≤60 months^20^, and median of 27.6 months^19^). In one study, 66.7% received tofacitinib and 86.7% were on background immunosuppressive therapy.^20^ For the other study, 72.7% of patients received background immunosuppressive treatment.^19^

Integration of these two datasets identified ten distinct cell populations, annotated with marker genes (91,196 cells), which captured the known cell populations in the skin (**Figure 3B**, **S5A)**. PDE4B expression was found to be most prominent and increased in SSc in both myeloid and lymphoid immune cells, as well as eccrine cells (**Figure 3C-D).** Notably, other SSc-relevant cell types including endothelial cells, fibroblasts, and pericytes showed low PDE4B expression overall and similar expression patterns between SSc and HCs (**Figure 3C-D**, **S5B)**. When PDE4B expression was compared at the individual patient level, eccrine cells demonstrated a significant increase in SSc compared to HCs (**Figure S5B**). Furthermore, endothelial cells also showed a significant increase in PDE4B expression in SSc, but a near-zero expression level. Other cell types, including fibroblasts and pericytes, showed low PDE4B expression overall and no significant upregulation in SSc (**Figure S5B**).

Given the observed increase of average cell-level PDE4B expression in myeloid cells alongside their known role in SSc pathogenesis, we analysed them further. Samples with fewer than 100 myeloid cells lacked sufficient statistical power for downstream analysis, resulting in a smaller number of HCs (2 HC, 19 SSc). The myeloid cells were annotated into nine populations (**Figure 3E**). Average PDE4B expression was observed to be increased in SSc across most myeloid cell populations (**Figure 3G)**. The strongest increase in PDE4B expression in SSc patients compared to HCs was observed in the Myel1 tissue-resident macrophage population (TRM, **Figure 3G**). While plasmacytoid dendritic cells (pDCs) were absent in HCs, the PDE4B expression levels in SSc patients were relatively high **(Figure 3G)**.

IHC staining of SSc and HC skin biopsies showed an overall increase in PDE4B^+^ cells in SSc consistent with the bulk RNA-seq data (p = 0.073, **Figure 4A-C**). PDE4B protein expression was predominantly localised to the epidermis, in perivascular structures and around eccrine structures (**Figure 4A-C**). Multiplex IF stainings confirmed that PDE4B was perivascularly expressed in SSc skin. However, PDE4B was not expressed in the vascular structures themselves as shown by limited co-localization with CD31 (mean expression 1.65% in SSc, 2.73% in HCs) and von Willebrand Factor (vWF, mean expression 1.80% in SSc and 1.23% in HCs, **Figure 4D, Figure S6A-E**). Next, we quantified PDE4B^+^CD68^+^ double-positive macrophages in SSc skin, where the percentage of double-positive cells was heterogeneous across patients, reflecting the observed transcriptional pattern (**Figure S6A, 3G**).

**Figure 4.**
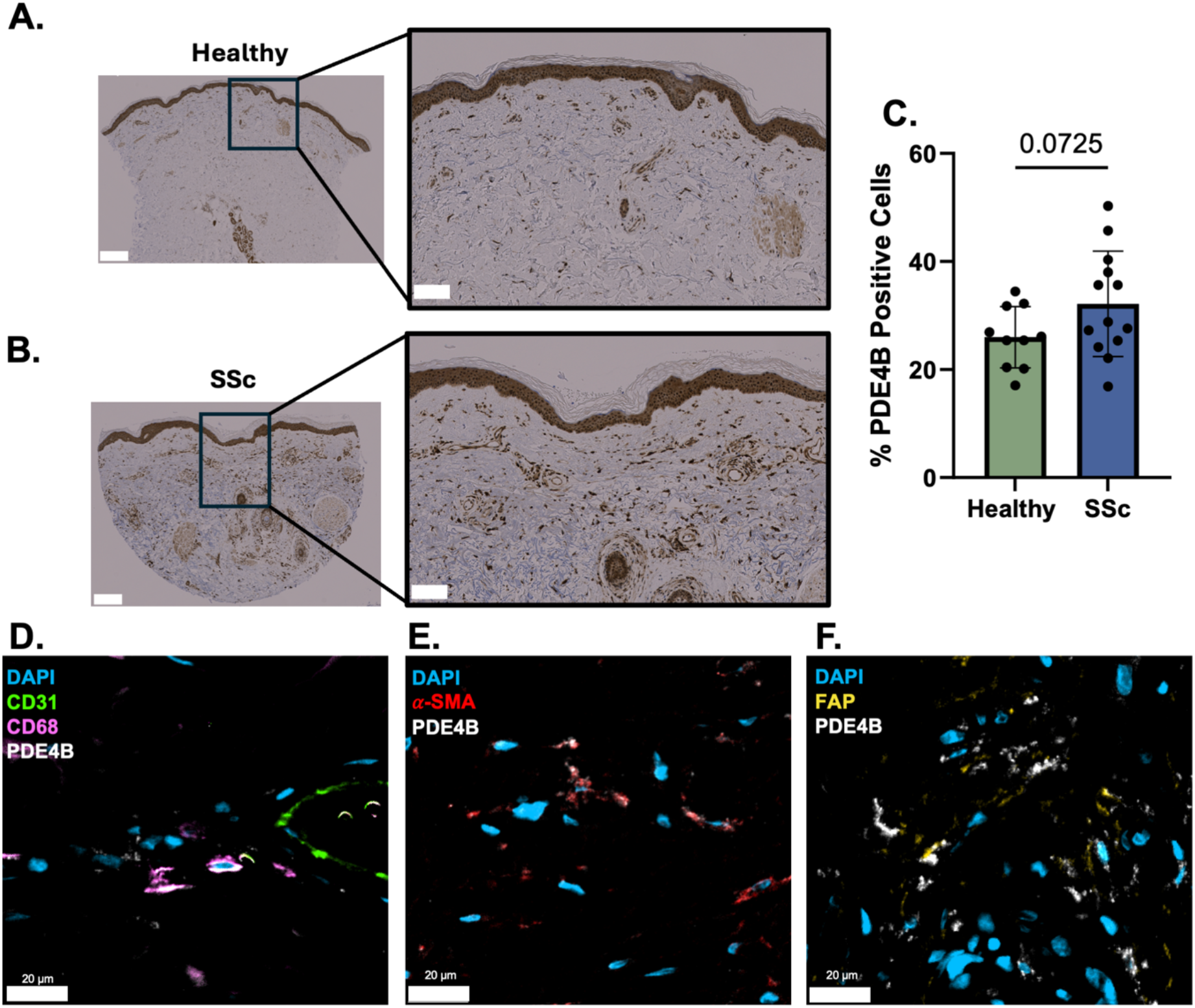
Immunofluorescent stainings show an increased amount of PDE4B^+^FAP^+^ cells in SSc skin. **(A)** Representative immunohistochemical staining of PDE4B protein in HC skin. Left panel at 2x magnification (scale bar = 200 μm); right panel shows magnified region at 5x magnification (scale bar = 100 μm). **(B)** Representative immunohistochemical staining of PDE4B protein in SSc skin. Left panel at 2x magnification (scale bar = 200 μm); right panel shows magnified region at 5x magnification (scale bar = 100 μm). **(C)** Quantification of the percentage of PDE4B-positive cells in HC (n = 10) and SSc (n = 13) skin sections, performed using QuPath. A trend towards increased PDE4B expression was observed in SSc compared to HC (p = 0.0725, Welch’s t-test). **(D)** Multiplex IF staining of endothelial cells (CD31^+^), macrophages (CD68^+^) and PDE4B in SSc skin. Scale bar = 20 μm. **(E)** Multiplex IF staining of activated/myofibroblasts (alpha smooth muscle actin, α-SMA^+^) and PDE4B in SSc skin. Scale bar = 20 μm. **(F)** Multiplex IF staining of activated fibroblasts (fibroblast activation protein, FAP^+^) and PDE4B in SSc skin. Scale bar = 20 μm. Abbreviations: HC, healthy controls; SSc, systemic sclerosis; IF, Immunofluorescent.

Given the proposed anti-fibrotic mechanism of PDE4B inhibitors, our multiplex IF panel included markers of activated fibroblasts. This found a co-expression of PDE4B in both α-smooth muscle actin (α-SMA) and fibroblast activation protein (FAP)^+^ cells in SSc skin (**Figure 4E**, **F**). An increased percentage of α-SMA^+^PDE4B^+^ cells was observed, with 94.7% of α-SMA^+^ cells expressing PDE4B (**Figure S6A**). Concurrently, the percentage of FAP^+^PDE4B^+^ cells was also elevated in SSc compared to HCs, confirming an increased PDE4B expression across activated fibroblasts (**Figure S6A**).

To this end, we quantified FAP^+^PDE4B^+^ cells from additional IF stainings, which revealed a significant increase in SSc skin compared to HCs (**Figure 5A-D**). As expected, FAP^+^ fibroblasts were significantly higher in SSc skin (17.4% of total detections) compared to HCs (7.26%). When calculating the proportion of FAP^+^ cells that also expressed PDE4B, approximately 90% of FAP^+^ cells in SSc skin expressed PDE4B compared to 19% in HCs (**Figure 5D**).

**Figure 5.**
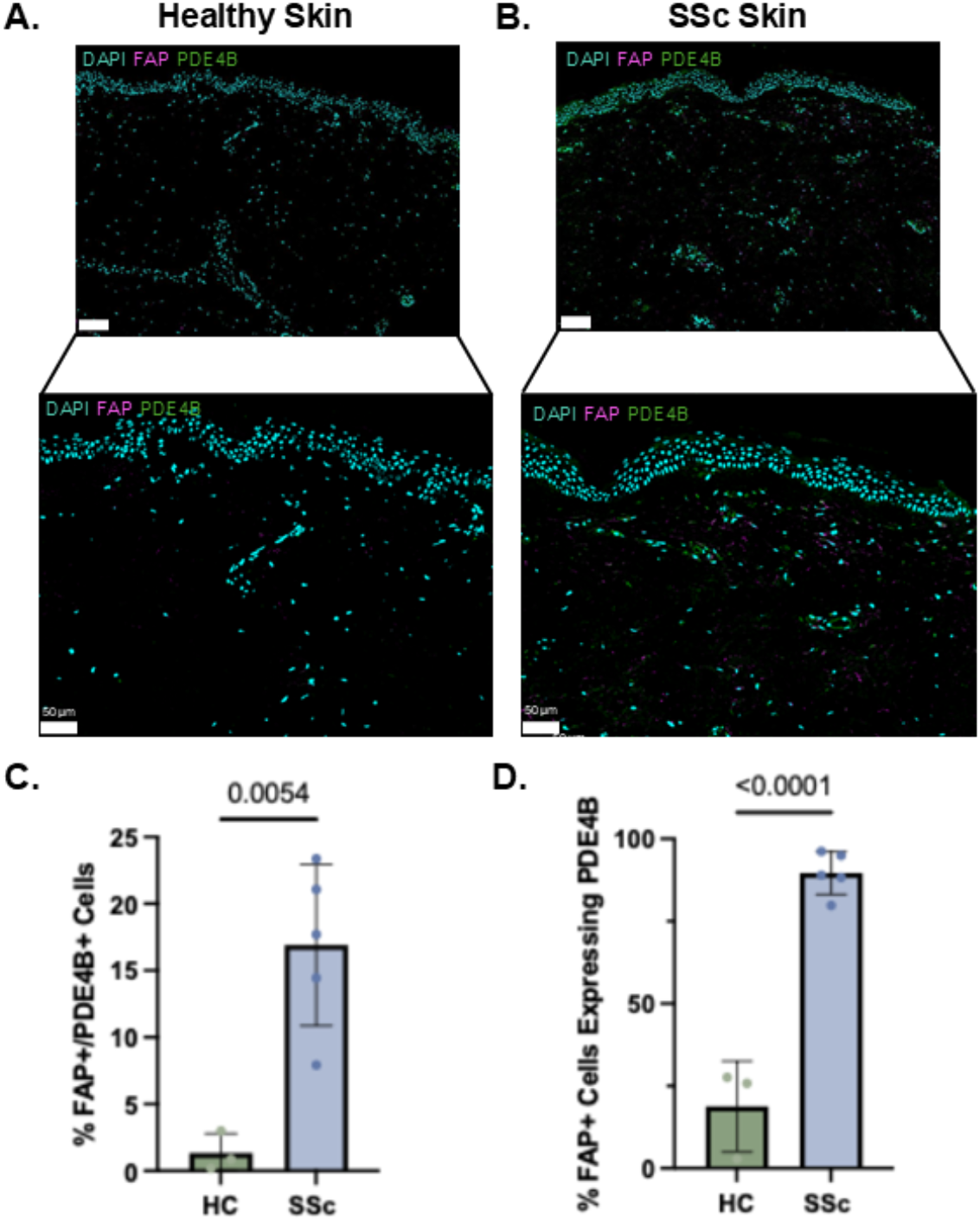
SSc skin shows a significant increase of PDE4B^+^FAP^+^ cells. **(A)** Co-staining of Fibroblast Activation Protein (FAP) and PDE4B in HC skin and **(B)** SSc skin. **(C)** Quantification of PDE4B^+^FAP^+^ double positive cells relative to the total number of cells detected, with each dot being a biological replicate (n = 3-5), p-value calculated using an unpaired t-test. **(D)** Quantification of FAP^+^ cells co-expressing PDE4B relative to the total number of FAP^+^ cells, with each dot being a biological replicate (n = 3-5), p-value calculated using an unpaired t-test. Abbreviations: HC, healthy controls; SSc, systemic sclerosis.

## Discussion

Despite the clinical momentum surrounding PDE4B inhibition, expression of PDE4B in SSc tissues and circulating cells was not systematically characterized, leaving a central piece of the target rationale unaddressed. In the present study, analysing key fibrotic organs and PBMCs PDE4B was consistently detected in immune cell populations, identifying the immune compartment as the most reproducible site of PDE4B expression in SSc. However, the cellular distribution and direction of regulation differed by tissue, reflecting compartment-specific aspects of PDE4B expression in SSc.

Monocytes and macrophages are key cells in the innate immune system, with critical roles in SSc.^24–26^. Macrophage PDE4B expression was decreased in both SSc-ILD and IPF lungs compared to HCs. This decrease in PDE4B expression was primarily driven by the shift in expression in FCN1^hi^ and FCN1^hi^ IFN-responsive populations, which are monocyte-derived macrophages that are recruited to sites of inflammation or injury.^23^ This may reflect a compositional shift in the fibrotic lung macrophage pool, in favour of tissue-resident phenotypes that intrinsically express less PDE4B. However, in IPF lungs, the FABP4^hi^ and a mixed population of FABP4^hi^/SPP1^hi^ macrophages showed increased PDE4B expression, indicating that tissue-resident alveolar macrophages display PDE4B expression dysregulation in IPF but not SSc-ILD. In the periphery, intermediate monocytes were the subset with the strongest PDE4B increase in SSc. Intermediate monocytes secrete TNF in response to LPS stimulation and have been identified to contribute to an aberrant IFN-response in SSc.^26,27^ Treatment of PBMCs with nerandomilast reduced TNF-α production.^17^ Taken together with our findings, these observations suggest that intermediate monocytes could be key mediators of the immunomodulatory effects associated with PDE4B inhibition. In SSc skin, PDE4B was most prominently expressed in the myeloid compartment, and specifically in tissue-resident macrophages. Overall, PDE4B expression in the monocyte/macrophage lineage was the most consistently altered feature across tissues and compartments in SSc.

T-cells have also been implicated in SSc pathophysiology and disease progression, with activated T-cells showing a bias toward a T helper 2 (Th2) phenotype.^25^ Following activation through the T-cell receptor (TCR), T-cells exhibit a transient increase in intracellular cAMP levels, which serves as a negative regulator of TCR signaling and effector function.^26^ This signaling is tightly controlled by PDEs.^26^ CD8⁺ T-cells emerged as a cross-compartmental source of PDE4B dysregulation in SSc, with consistent upregulation observed in both lung and PBMCs. The PDE4B2 isoform regulates CD8⁺ T-cell function by degrading cAMP, thereby relieving its inhibitory effect and promoting IL-2 signaling and TCR activation.^26^ cAMP is a molecular brake for TCR signaling and effector molecule production. Thus, elevated PDE4B expression in SSc-ILD CD8⁺ T-cells is consistent with an activated effector state, in which this regulatory brake is diminished and may be restored through PDE4B inhibition.^27^ Importantly, the SSc-ILD-specific increase in CD8⁺ T-cells distinguishes the SSc immune signature from IPF and may represent a disease-specific target for therapeutic intervention. In SSc-ILD, the CD4⁺ITGB1⁺ T-cell subset showed increased PDE4B expression. ITGB1 (encoding CD29) marks an activated, cytotoxic memory CD4⁺ T-cell subset whose effector activity has been implicated in driving persistent tissue inflammation in chronic immune-mediated disease.^28,29^ PDE4B expression was unchanged in CD4⁺ T-cells in SSc PBMCs and skin. These findings indicate that CD4⁺ PDE4B dysregulation in SSc-ILD is restricted to an activated memory subset, rather than representing a global change in the CD4⁺ effector compartment.

Within the PBMCs, we aimed to address the heterogeneity of PDE4B expression across patients. This presented in a pattern that could be classified into three distinct patient endotypes: PDE4Bˡᵒ, PDE4Bᵐᵉᵈ, and PDE4B^hi^. This stratification was consistent within patients across the major PBMC cell types. PDE4B^hi^ patients could not be distinguished using standard clinical variables, indicating that direct measurement of PDE4B expression is required to stratify patients for clinical trial inclusion or targeted treatment.

Fibroblasts are the principal effector cells of fibrosis, driving extracellular matrix deposition in fibrotic tissues, and direct anti-fibrotic activity through cAMP accumulation is a central component of the rationale for PDE4B inhibition.^29^ Transcriptionally, PDE4B was sparsely detected in fibroblasts in both skin and lung, with no significant difference between SSc and HCs. PDE4B mRNA may be underestimated in droplet-based single-cell data due to technical dropout and post-transcriptional regulation, limiting interpretation in certain cell types. On the protein level, PDE4B was highly expressed in FAP⁺ fibroblasts and α-SMA⁺ myofibroblasts. FAP is largely absent from healthy adult tissue but is selectively upregulated on activated fibroblasts, with the FAP⁺ population in SSc skin representing a pathologically expanded fibroblast pool central to fibrosis.^34^ These findings suggest that PDE4B upregulation is not simply driven by increased FAP⁺ cell abundance but instead reflects a selective and near-universal enrichment of PDE4B within the activated fibroblast compartment in SSc skin. This provides a clear rationale at the protein level for the direct anti-fibrotic effects of PDE4B inhibition in the skin reported *in vitro* and *in vivo*.^16,17^ Although PDE4B inhibition is proposed to promote endothelial stabilization, no transcript-or protein-level dysregulation was observed in SSc. Therefore, this study does not provide evidence to support this rationale.

Several limitations should be acknowledged. First, our analysis assessed PDE4B expression at the transcript and protein levels but did not assess enzyme activity, post-translational regulation, or isoform composition. The relationship between PDE4B expression and functional cAMP hydrolysis cannot be inferred from expression data alone. Future studies should couple expression measurements with direct PDE4B activity assays in primary SSc cells or advanced *in vitro* models to link the patterns described here to the functional consequences of PDE4B inhibition. Furthermore, protein level confirmation was only available for SSc skin, but the transcript–protein discordance observed in fibroblasts underscores the value of this validation and warrants extension to lung tissue. Additionally, the cross-tissue patterns reported here derive from independent patient cohorts rather than paired samples, with different background treatment in the lung, PBMC, and skin datasets. Notably, earlier nerandomilast trials excluded pulmonary specific background immunosuppressive therapy, whereas the ongoing and upcoming trials permit it. This raises the concern that concurrent immunosuppression could attenuate the effect of nerandomilast. However, many patients in our datasets were receiving immunosuppressive therapy at the time of sampling and still showed elevated PDE4B expression, indicating that immunosuppression does not abolish the PDE4B signal.

## Conclusion

Together, these findings provide the first cell-type-resolved characterization of PDE4B in SSc, demonstrating consistent immune-cell dysregulation across the lung, PBMCs, and skin. Furthermore, the selective enrichment of PDE4B at the protein level within activated fibroblasts, together with marked patient-level heterogeneity, supports PDE4B as a disease-relevant target in SSc.

## Methods

### Cohort Description & Ethical Statement

The collection and usage of human samples used to generate in-house datasets was approved by the Cantonal Ethics Commission (KEK) Zurich (KEK-Nr. 2018-01873, KEK-ZH 515, PB-2016-02014). Patients and the public were not involved in the design, conduct, reporting, or dissemination plans of this research. Informed consent was provided by all patients and healthy controls (HCs). All research followed the Declaration of Helsinki for research ethics. All patient material included in in-house generated datasets was obtained from patients fulfilling the ACR/EULAR 2013 criteria for SSc. Other datasets used in this study are described in **Table 1**. Detailed information regarding patient demographics and data analysis can be found in **Supplemental Files 1-4.**

### Immunohistochemical (IHC) & Immunofluorescent (IF) Stainings

Skin samples (3 µm sections) were stained for PDE4B (Novus, #NBP2-01171) on the BOND-MAX (Leica) with hematoxylin counterstain (reagents in **Supplemental File 4**). Whole-slide scans were acquired at 20x on the Axioscan.Z1 (Zeiss), and the percentage of PDE4B-positive cells was quantified in QuPath using positive cell detection.^31^ For IF stainings, whole-section images were acquired on the Leica CrestOptics CICERO spinning disk and analyzed in QuPath. Detailed methodological descriptions are provided in **Supplemental Files 1** and **4**. Statistics were performed in GraphPad Prism 10.6.1.

### Multiplex IF Stainings

High-plex spatial protein profiling was performed on 4 µm skin sections using the COMET™ platform (Lunaphore, antibodies and reagents in **Supplemental File 4**). Images were processed in the Horizon™ viewer, and positive cells were quantified in QuPath. Detailed information can be found in **Supplemental File 1.**

### Lung single-cell RNA sequencing

Publicly available scRNA-seq datasets from the national center for biotechnology information (NCBI) ascension numbers GSE128169 and GSE128033 were used. We downloaded Cell Ranger–processed gene expression matrices (v2.0) from GEO. Base calling, alignment, and generation of gene–barcode matrices were performed by the original study with the GRCh38 reference genome. These two datasets were integrated, resulting in one analysis with SSc– associated interstitial lung disease (SSc-ILD, n = 4, GSE128169) samples, idiopathic pulmonary fibrosis (IPF, n = 4, GSE128033) samples, and healthy control (HC) lung (n=4) samples (n = 4, GSE128169). For certain patients, biopsies for both the upper and lower lobes were taken. In these cases, the expression values were aggregated within each individual to obtain a per-patient measurement prior to any downstream statistical analysis. The analysis was performed using R version 4.4.2 and Seurat version 5.2.1.

For quality control, first doublets were removed using scDblFinder and an initial filtering of cells with < 200 counts was performed. Subsequently, additional filtering was performed. In short, the 10^th^ and 90^th^ percentiles of each quality control parameter (nFeature, nCount, and mitochondrial content) were used in addition to cells having to have >200 genes, >500 UMIs, <15% mitochondrial content, and <30% ribosomal content. Remaining cells were normalized (log-transformed normalized expression variables) and the top 2000 variable features were selected. Data scaling and PCA dimensionality reduction (number of PCs were manually selected using ElbowPlot) were done using Seurat. To correct batch effects and inter-patient variability, Harmony (v1.2.3) integration was used.

Major cell types were identified, with neighbours, clusters and Uniform Manifold Approximation and Project (UMAP) calculated using Seurat. Major cell types were manually annotated using the top 10 markers for each cluster. For annotation of subpopulations within the cell types, cell types of interest were selected and reclustered, using STACAS (v2.2.2) for integration. The same workflow as detailed above was followed.

Differential expression analysis was performed using the FindMarkers function, implementing the MAST test. To adjust for inter-sample variability, each sample was used as a covariate. To define significant cell types, a |log2FC| > 0.5 was used as a cut-off.

### PBMC Isolation & scRNA-seq

Whole blood was taken in BD Vacutainer^®^ EDTA tubes (CAT #367863) from SSc patients and healthy controls. Peripheral blood mononuclear cells (PBMCs) were isolated from whole blood using density gradient centrifugation using LymphoPrep density gradient medium (STEMCELL Technologies, #18060) and SepMate™ tubes (STEMCELL Technologies, #85450). 500,000 cells from each donor (25 SSc patients, 9 healthy controls) were stained with a panel of ADT antibodies for 30 minutes at 4°C. After the staining, cells were washed and strained to ensure single cell suspension. Then, cells were re-counted and diluted as necessary for appropriate input into the 10X Chromium chip (10X Genomics). Two samples were pooled together per single cell reaction and each sample labeled with a hashtag antibody for Beta-2-microglobulin (Biolegend). Single cells and barcoded gel-beads in emulsion were co-encapsulated in droplets using the Chromium Single Cell Controller Instrument (10X Genomics) to allow RNA barcoding and retrotranscription of each single cell. Libraries for surface markers and cDNA tags were prepared. Single cell preparation procedures were performed following the protocol for Chromium Next GEM single cell 3’ reagent kit v3.1 with feature barcoding technology for cell surface proteins. Library size and concentration were assessed using the Agilent Bioanalyzer (Agilent Technologies). Sequencing was performed at the Functional Genomics Centre Zurich (FGCZ) using Illumina NovaSeq 6000. Six thousand cells per patient were used and 50,000 reads as mean read depth per cell were targeted. Clinical and demographic data were prospectively collected as part of the local registry following EUSTAR standards and definitions. Clinical data were available for all patients, including disease duration, auto-antibody status, skin involvement (diffuse vs limited cutaneous/sine scleroderma SSc), r-EUSTAR-AI, CRP, ESR, mRSS, FVC, DLCO, KCO, disease duration. Patient demographics can be found in **Table 2**.

Reads were aligned using CellRanger (*CellRanger mkfastq* (v6.1.2)) and Seurat (v4.3.0) was used for analysis of all major cell types, whereas Seurat (v.5.2.1) was used for detailed analyses. Data-preprocessing consisted of the following: demultiplexing of samples, doublet detection using scDblFinder, and cells were filtered based on the number of detected genes (nFeature_RNA) and mitochondrial gene content (percent.mt). PBMCs with fewer than 200 or more than 6,000 detected genes, and cells with >15% mitochondrial reads, were excluded from downstream analyses. Subsequently, normalisation (*scTransform),* and identification of the top 2,000 highly variable genes was performed. The data was then scaled and PCA dimensionality reduction was performed with manual PC selection using the ElbowPlot function in Seurat. Major cell types were annotated using top markers genes (FindAllMarkers function in Seurat) and confirmed by automatic annotation using the Azimuth PBMC reference (https://azimuth.hubmapconsortium.org).^35^ Antibody-derived tag (ADT) counts were normalised using centered log-ratio (CLR) transformation to account for technical variation in antibody capture efficiency and sequencing depth across cells. Differential gene expression analysis was performed for each major PBMC population (Monocyte, dendritic cells, NK cells, B cells, CD4^+^ T-cells, CD8^+^ T-cells) using the FindMarkers function in *Seurat* with the MAST framework using sex as a co-variate.

Raw pseudobulk count matrices were generated using the AggregateExpression function from the Seurat package. Counts were normalised using the edgeR package by constructing a DGEList object and applying the trimmed mean of M-values (TMM) normalization method to account for differences in library size and composition. Normalised expression values were then transformed to log2 counts per million (log2-CPM) using a prior count of 1 to stabilise variance. The resulting log2-CPM expression matrix was used for all downstream analyses. Using the normalised pseudobulk PDE4B expression values, the mean and standard deviation of the healthy controls group were calculated. PDE4B expression in each sample was then standardised by computing a Z-score relative to the HC distribution

### .Bulk RNA-seq data

Bulk RNA-seq data from the Prospective Registry for Early Systemic Sclerosis (PRESS) was used. In short, Raw reads were mapped to the human genome hg38, downloaded from the University of California Santa Cruz Genome Bioinformatics site (http://genome.ucsc.edu), with no more than two mismatches for each read, using TopHat v2.1.11.2. Subsequently, transcript count value was obtained using htseq-count,4 using default parameters. Further information can be found in the original study.

### Skin single-cell RNA sequencing

Each dataset was preprocessed independently using Scanpy (v1.11).^36^ Quality control was performed by applying thresholds on the following per-cell metrics: mitochondrial RNA fraction (< 20%), ribosomal RNA fraction (> 5% and < 45%), haemoglobin and platelet-associated transcript fraction (< 0.5%), total UMI count (< 40,000), and number of detected genes (> 500 and < 5,000). Doublets were identified and removed using Scrublet.^37^ Samples retaining fewer than 500 cells after filtering were excluded from downstream analyses. Gene expression counts were normalised to 10,000 counts per cell followed by log1p transformation. Highly variable genes were selected, principal component analysis (PCA) was performed on the scaled data, and a neighbourhood graph was constructed using n_neighbors = 15 and n_pcs = 9, followed by UMAP dimensionality reduction.

Prior to integration, the comparability of normalised expression distributions across datasets was systematically evaluated. The mean and standard deviation of normalised log-transformed counts were computed for each dataset and compared through density plots and quantile–quantile (Q-Q) plots to assess distributional similarity. Potential residual batch effects were further characterized by examining differences in per-cell gene count and UMI count distributions across samples, as well as by assessing the expression of established cell-type marker genes across datasets. These analyses confirmed sufficient cross-dataset comparability as a prerequisite for multi-dataset integration.

Cell-type annotation was performed using CellTypist (v1.7.1) by applying the Adult_Human_Skin model.^35^ Automated annotations were subsequently reviewed and manually curated to resolve ambiguous or inconsistent assignments and to harmonise nomenclature across tissue compartments. In the differential expression analysis, it was not possible to evaluate differences in cell populations that had fewer than 100 cells per sample, leading to sample drop-out in certain conditions.

### Immunohistochemistry (IHC) Staining of PDE4B in SSc Skin

For IHC of skin biopsies, 3 µm skin biopsies were fixed in 4% paraformaldehyde (FormaFix, #2311158) for 12-16 hours and subsequently transferred to 50% ethanol. Tissue processing was performed using the HistoCore PELORIS 2 system (Leica Biosystems) according to standard protocols. After tissue processing, skin biopsies were embedded in paraffin using the HistoCore Arcadia H Embedding Station (Leica Biosystems). 3 µm sections were cut using the HistoCore Autocut microtome (Leica Biosystems), transferred on Superfrost Plus slides (VWR, #631-0446) and dried overnight at 58°C before deparaffinisation in the BOND-MAX using the Bond Dewax Solution (Leica Biosystems, AR9222).

Histological staining with hematoxylin and eosin (H&E) was performed to assess tissue morphology according to standard protocols. IHC was performed for PDE4B (Novus Biologics, #NBP2-01171) on the BOND-MAX (Leica Biosystems) using the BOND Polymer Refine Detection Kit (Leica Biosystems) according to the manufacturer’s instructions. In short, samples were deparaffinised using the Dewax Solution (Biosystems, AR9222). Subsequently, the deparaffinised sections were subjected to heat-induced epitope retrieval (HIER) using ethylenediaminetetraacetic acid (ETDA)-based buffer (pH 9, BOND Epitope Retrieval Solution 2, Leica Biosystems) for 20 minutes at 95°C. Subsequently, non-specific antibody binding was blocked, and sections were incubated with the primary and secondary antibodies. Lastly, 3,3’-Diaminobenzidine (DAB) was used for the antibody detection chromogen and sections were counterstained with hematoxylin. Detailed information regarding reagents and antibodies can be found in **Supplemental Table 4.**

Whole slide scans of IHC stained skin sections were imaged at a 20X magnification on the Axioscan.Z1 (Zeiss) slide scanner at the Centre for Microscopy and Image Analysis. Quantification of PDE4B expression in skin was performed using QuPath (0.4.3) to measure the percentage of positively stained cells. In short, the “positive cell detection” feature was used in QuPath (v.6) by detecting nuclei using the hematoxlin optical density (OD). Subsequently, DAB was used to identify stained cells (cell threshold DAB OD mean). Statistical analysis was performed in Graphpad prism version 10.6.1.

### Immunofluorescent Stainings

Immunofluorescent stainings were performed on SSc and healthy skin samples processed as previously described. Sections were deparaffinised and underwent heat-induced epitope retrieval in citrate buffer (pH6). Subsequently, sections were washed in TBS-Tween and blocked with 10% horse serum (S-2000) in antibody diluent (Dako, S3022) for 1 hour. Sections were incubated with primary antibodies in antibody diluent or in only antibody diluent for secondary controls overnight. Sections were then washed with 3% milk (Fisher Scientific, #232100) in TBS-Tween and incubated with the secondary antibody for 1 hour in the dark. Subsequently, autofluorescence was reduced (Vector® TrueVIEW® Autofluorescence quenching kit, Vector Laboratories, SP-8400-15) and samples were incubated with DAPI for 15 minutes, after which the slides were mounted (Vector Laboratories, H-1700-10). Detailed information regarding reagents and antibodies can be found in **Supplemental Table 4.**

Imaging was performed on the Leica CrestOptics CICERO Spinning Disk at the Centre for Microscopy and Image Analysis, and whole section images were acquired. Images were processed by manually adjusting the brightness and contrast for visualisation in QuPath v0.7.0, after which analysis of double cell detection was performed using positive cell detection for nuclei and each staining, which were subsequently combined into a single threshold classifier.

### Multi-Plex Immunofluorescent Stainings

High-plex spatial protein profiling was performed on FFPE skin tissue sections using the COMET™ platform (Lunaphore). Skin samples (4 µm) were sectioned transferred onto Superfrost Plus slides (VWR, #631-0446), baked at 60 °C for 1 hour, and subsequently underwent manual deparaffinisation with xylene (2 times 5 minutes), and 100% ethanol (2 times 2 minutes). Sections were then dried at 60 °C for 5 minutes. Heat-induced epitope retrieval was formed using the PT module (Epredia) at 99 °C for 1 hour using buffer H (pH 9 EDTA). Finally, blocking in 5% horse serum (Vector Laboratories, S2000-20) was performed for 30 minutes.

Sequential immunofluorescence (SeqIF™) was performed for a custom panel of antibodies across 13 cycles of primary antibody incubation, secondary antibody detection, imaging, and finally, elution of the antibodies prior to the next cycle. Signal detection was performed for TRITC-and Cy5-conjugated antibodies. Detailed information regarding reagents and antibodies can be found in **Supplemental Table 4.** After the image acquisition, images were processed using the Horizon™ viewer software (Version 2.5.5.1). Here, background subtraction was performed and the brightness of stainings manually adjusted per image. Quantification was performed in QuPath (0.7.0). Nuclei were segmented using the InstanSeg plugin (0.1.7) and cell boundaries were approximated by expanding nuclear masks by 3µm. Incorrectly segmented objects were excluded using a random forest object classifier. Cells positive for each marker were identified by thresholding mean per-cell intensity, with thresholds set manually and applied consistently across all samples.

### Generative AI Statement

During the preparation of this work, the authors used Claude (Anthropic) and Microsoft Copilot in order to rephrase text for clarity. After using these tools, the authors reviewed and edited the content as needed and take full responsibility for the content of the published article.

## Acknowledgements

We would like to thank the Center for Microscopy and Image Analysis for the use of their microscopes and for support during image processing and analysis. We gratefully acknowledge the PRESS investigators for sample collection and data access, and the Functional Genomic Center Zürich for technical support and expertise. We also thank Peter Künzler and the other members of the laboratory for helpful discussions and technical assistance. Finally, we wish to thank all the patients who contributed to this research and made this work possible.

## CRediT Authorship Contributions

**Astrid Hofman –** Conceptualization, Data Curation, Formal analysis, Investigation, Methodology, Writing – Original Draft, Writing – Review and editing

**Pietro Bearzi-**Conceptualization, Data Curation, Formal analysis, Investigation, Methodology, Writing – Review and editing

**Sara Burckhardt -** Data Curation, Formal analysis, Methodology, Writing – review and editing

**Maryam Asadikorayam -** Formal analysis, Investigation, Methodology, Writing – review and editing

**Andrea Laimbacher -** Data Curation, Investigation, Methodology, Writing – review and editing

**Cristian Iperi-**Data Curation, Formal analysis, Investigation, Methodology, Writing – review and editing

**Muriel Elhai –** Investigation, Resources, Writing – review and editing

**Cosimo Bruni –** Data Curation, Formal analysis, Writing – review and editing

**Flurin Struzenegger -** Formal analysis, Methodology, Writing – review and editing

**Shervin Assassi -** Data Curation, Formal analysis, Investigation, Resources, Writing – review and editing

**Laura Much-** Data Curation, Investigation, Methodology, Writing – review and editing

**Anna-Maria Hoffmann-Vold –** Investigation, Supervision, Writing – review and editing

**Blaž Burja –** Formal Analysis, Investigation, Writing – review and editing

**Helen C. Jarnigan -** Data Curation, Formal analysis, Writing – review and editing

**Michael L. Whitfield -** Data Curation, Formal analysis, Writing – review and editing

**Mike O. Becker –** Resources, Writing – review and editing

**Lumeng Li –** Formal Analysis, Investigation, Writing – review and editing

**Philip Stauffer -** Formal Analysis, Investigation, Writing – review and editing

**Shihan Xu-** Formal Analysis, Investigation, Writing – review and editing

**Sophie Wagner -** Formal Analysis, Investigation, Writing – review and editing

**Yasmine Illi-** Formal Analysis, Investigation, Writing – review and editing

**Zhiyun Gong -** Data Curation, Formal analysis, Writing – review and editing

**Elena Pachera –** Conceptualization, Data Curation, Investigation, Supervision, Writing – Review and editing

**Oliver Distler –** Conceptualization, Data Curation, Investigation, Resources, Supervision, Writing – Review and editing

## Data Availibility

All data availability is indicated in the text, with references to the appropriate repositories/studies. The PBMC scRNA-sequencing data will be made publicly available upon publication.

---

**Figure S1.**
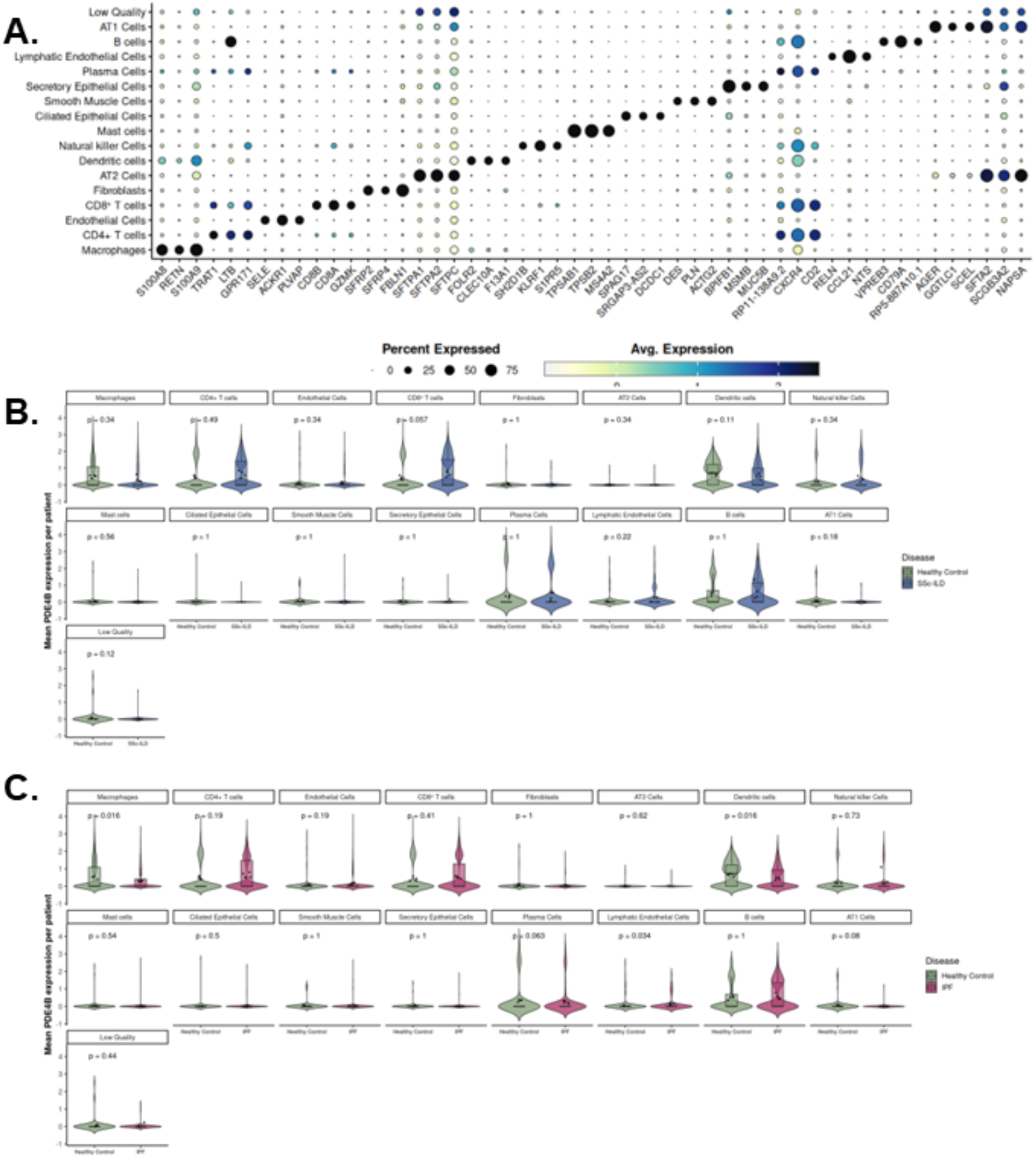
PDE4B expression across lung populations in SSc-ILD, IPF and HC lung. **(A)** Top 3 marker genes per cell population identified in the lung scRNA-seq. **(B)** Mean PDE4B expression per patient across lung cell types in HCs (n=4) and SSc-ILD (n=4). Statistical comparisons were performed using the Wilcoxon rank-sum test. Each dot represents one sample. **(C)** Mean PDE4B expression per patient across lung cell types in HCs (n=4) and IPF (n=4). Statistical comparisons were performed using the Wilcoxon rank-sum test. Each dot represents one sample. Abbreviations: SSc-ILD: systemic sclerosis-associated interstitial lung disease, IPF: Idiopathic pulmonary fibrosis, HC: healthy controls.

**Figure S2.**
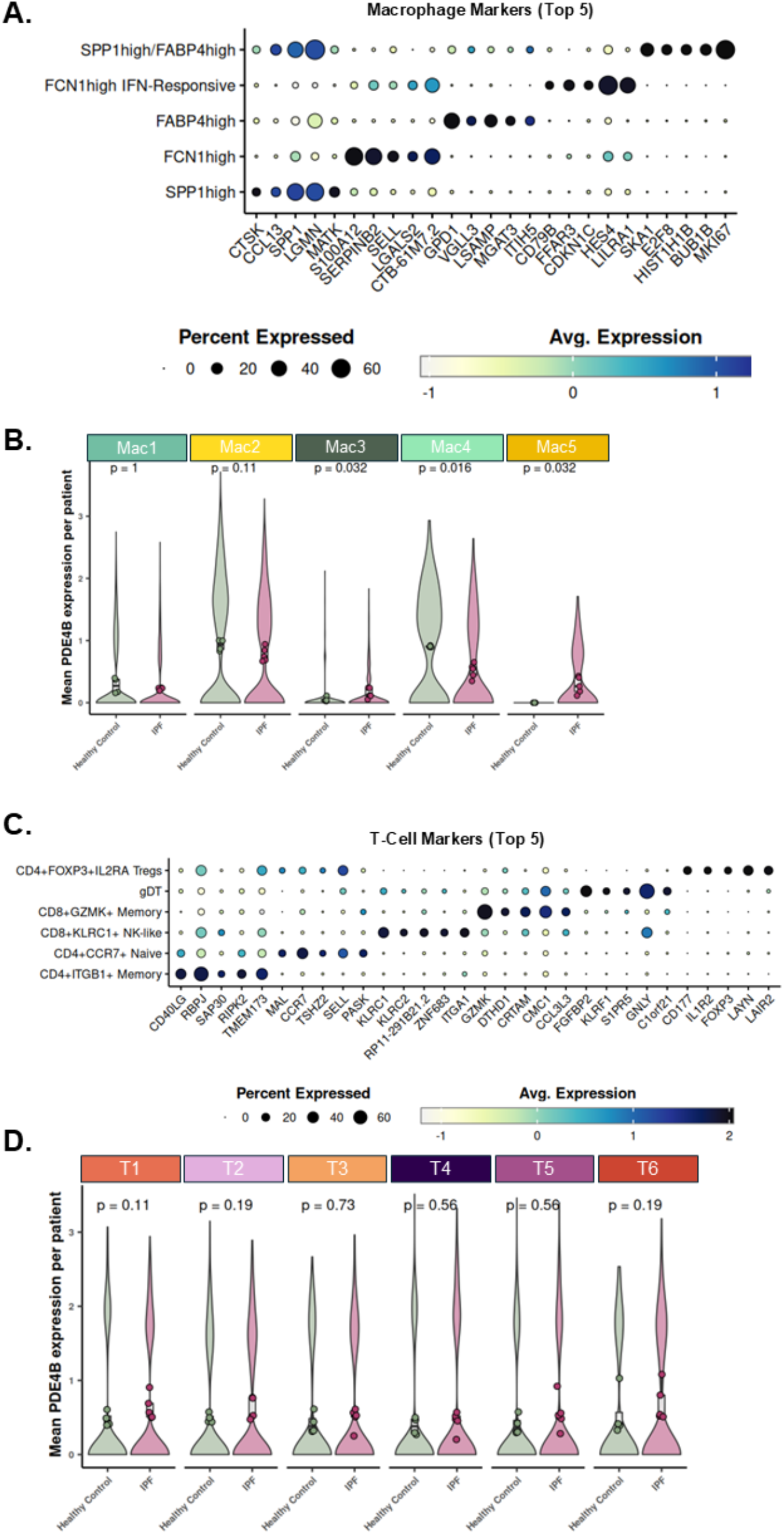
PDE4B Expression across macrophage and T-cell subpopulations in IPF and HCs. **(A)** Top 5 marker genes per macrophage population identified in the lung scRNA-seq. **(B)** Mean PDE4B expression per patient across macrophage populations in HCs (n=4) and IPF (n=4). Statistical comparisons were performed using the Wilcoxon rank-sum test. Each dot represents one sample. **(C)** Top 5 marker genes per T-cell population identified in the lung scRNA-seq. **(D)** Mean PDE4B expression per patient across T-cell populations in HCs (n=4) and IPF (n=4). Statistical comparisons were performed using the Wilcoxon rank-sum test. Each dot represents one sample. Abbreviations: SSc-ILD: systemic sclerosis-associated interstitial lung disease, IPF: Idiopathic pulmonary fibrosis, HC: healthy controls.

**Figure S3.**
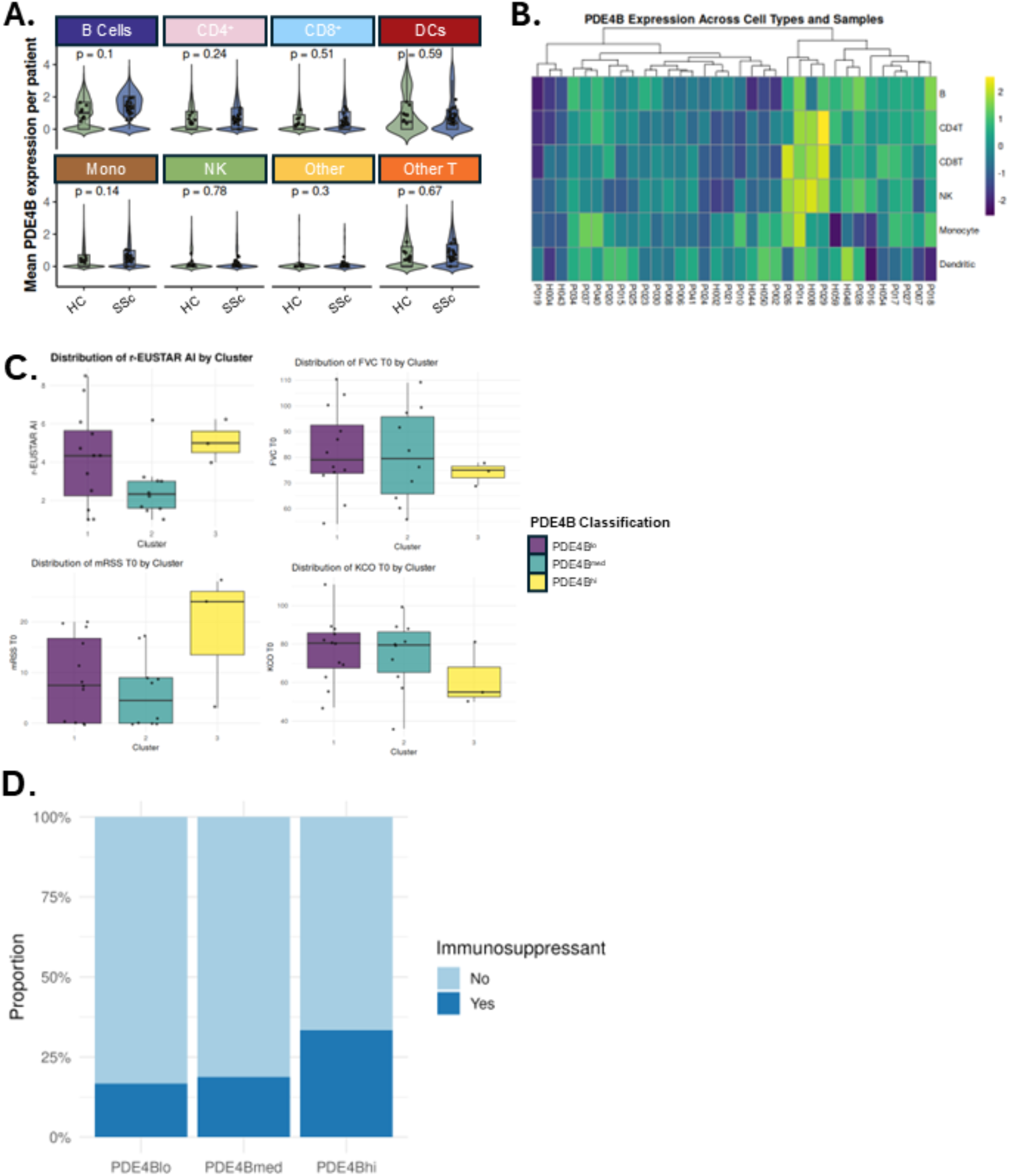
PDE4B expression in PBMCs in SSc patients and HCs. **(A)** Mean PDE4B expression per patient across PBMC populations in SSc (n=25) and HCs (n=9). Statistical comparisons were performed using the Wilcoxon rank-sum test. Each dot represents one sample. **(B)** Unsupervised clustering of Z-scores of PDE4B expression across SSc patients and HCs. **(C)** Boxplots of tested clinical variables across the different PDE4B expression patient endotypes. Each dot represents one sample. **(D)** Proportion of patients receiving immunosuppressive therapy across the PDE4B expression endotypes. Abbreviations: SSc: Systemic sclerosis, HC: healthy controls, PBMCs: peripheral blood mononuclear cells.

**Figure S4.**
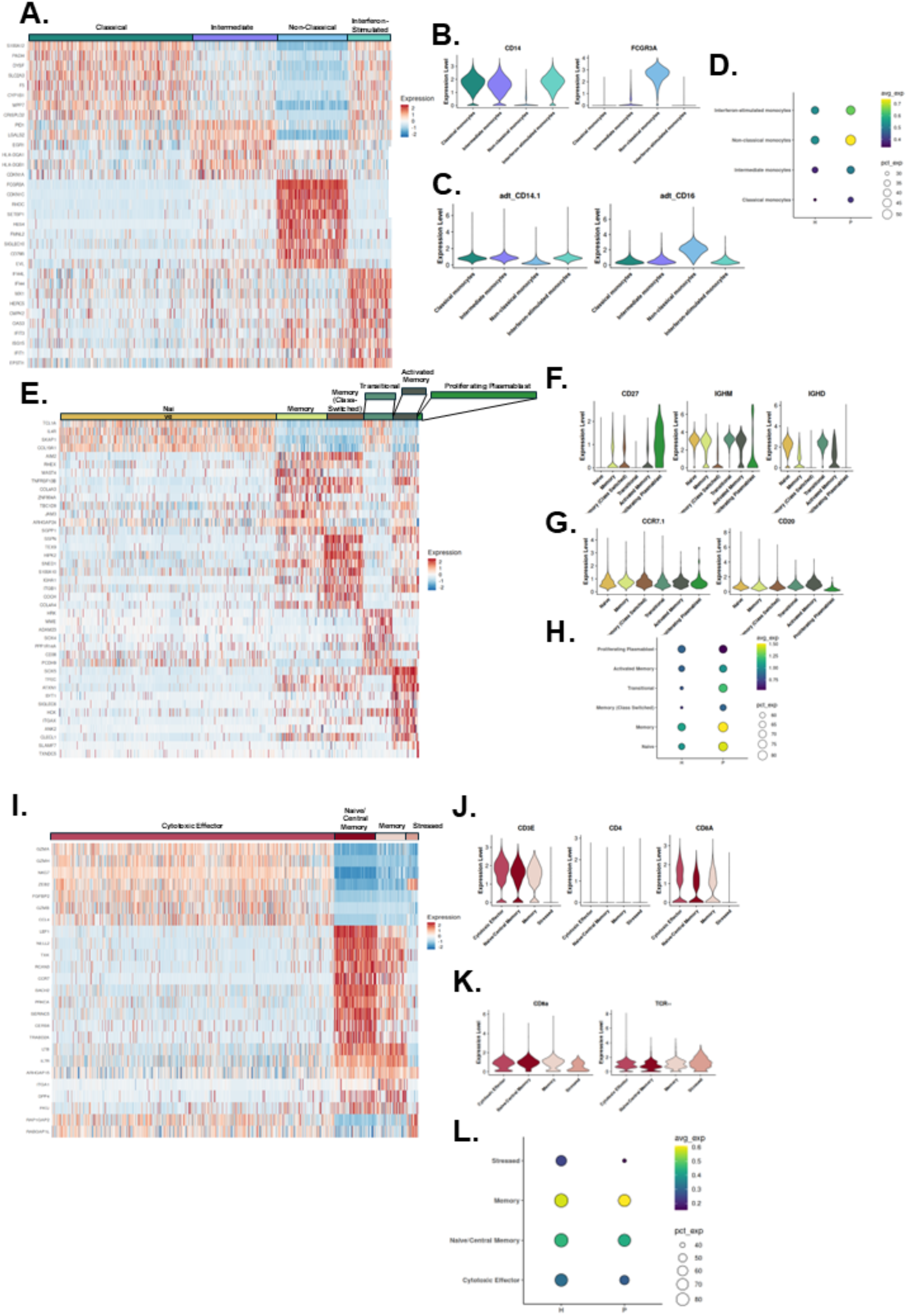
Marker genes in PBMC subpopulations and PDE4B expression across monocyte, B-cell and T-cell subpopulations. **(A)** Heatmap of scaled expression values for each monocyte subpopulation. **(B)** Violin plot of RNA markers for monocytes. **(C)** Violin plot of Cite-Seq markers for monocytes. **(D)** Dot plot showing the expression levels of PDE4B in each monocyte population, with dot size representing the percentage of expressing cells and colour indicating average expression level. **(E)** Heatmap of scaled expression values for each B-cell subpopulation. **(F)** Violin plot of RNA markers for B-cells. **(G)** Violin plot of ADT markers for B-cells. **(H)** Dot plot showing the expression levels of PDE4B in each B-cell population, with dot size representing the percentage of expressing cells and colour indicating average expression level. **(I)** Heatmap of scaled expression values for each T-cell subpopulation. **(J)** Violin plot of RNA markers for T-cells. **(K)** Violin plot of Cite-Seq markers for T-cells. **(L)** Dot plot showing the expression levels of PDE4B in each T-cell population, with dot size representing the percentage of expressing cells and colour indicating average expression level. Abbreviations: SSc: Systemic sclerosis, HC: healthy controls, PBMCs: peripheral blood mononuclear cells.

**Figure S5.**
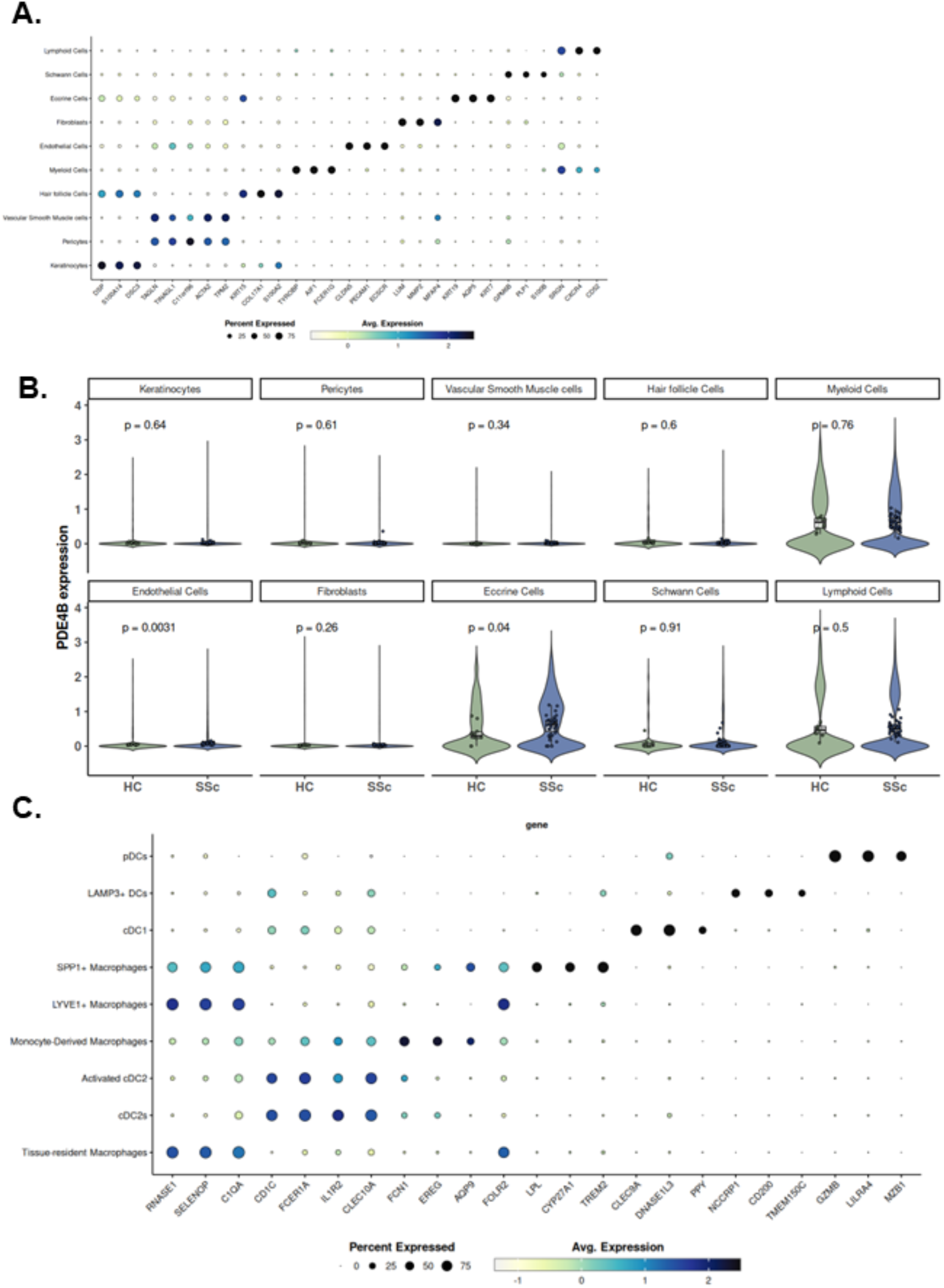
PDE4B expression in SSc skin. **(A)** Dot plot showing the top 3 markers for each annotated cell type in SSc skin. **(B)** Mean PDE4B expression shown for each cell type for SSc (n = 19) and HC (n = 10). Each dot represents one sample. **(C)** Dot plot showing the top 3 markers per cluster for the myeloid cell subpopulations. Abbreviations: SSc: Systemic sclerosis, HC: healthy controls.

**Figure S6.**
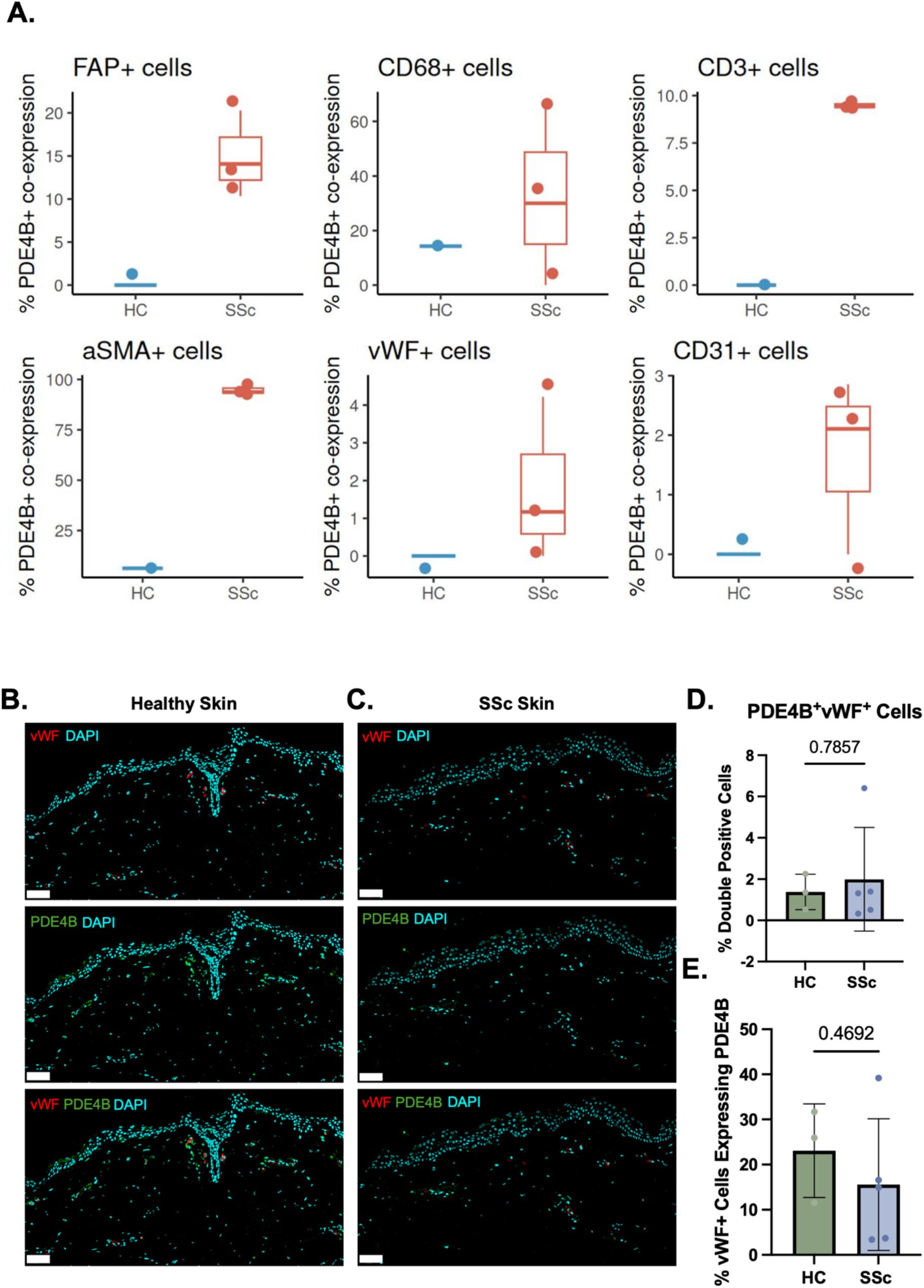
Overview of PDE4B expression using (multiplex) IF. **(A)** Quantification of Multiplex IF stainings for each marker of interest (n = 1-3) **(B)** Co-staining von Willebrand Factor (vWF) staining and PDE4B in healthy skin and **(C)** in SSc skin (scale bar 100 μm). **(D)** Quantification of vWF+PDE4B+ double positive cells, with each dot being a biological replicate (**n** = 3-5), p-value calculated using a Mann-Whitney t-test. **(E)** Quantification of vWF+ cells that also express PDE4B, with each dot being a biological replicate (n = 3-5), p-value calculated using a Mann-Whitney t-test. Scale bars: 100 µm. Abbreviations: SSc: Systemic sclerosis, HC: healthy controls, FAP: Fibroblast activation protein, aSMA: alpha-smooth muscle actin, vWF: von Willebrand Factor.

